# Elastase mediated white matter damage in cerebral small vessel disease: Microglia - neutrophils pas de deux

**DOI:** 10.1101/2024.12.23.630204

**Authors:** Cheng-Ya Dong, Weiqi Chen, Qi Li, Junwan Fan, Tian Lan, Wanlin Dong, Lina Sun, Yingjie Wang, Mingming Shi, Yongjin Huang, Yingying Chang, Ningning Wang, Jing Xue, Lingling Jiang, Yue Huang, Yuesong Pan, Wenyan He, Kaibin Shi, Xiaotong Ma, Yongjun Wang, Fu-Dong Shi, Alexei Verkhratsky, Yilong Wang, Wei-Na Jin

## Abstract

Cerebral small vessel disease (CSVD) leads to an extensive white matter damage associated with cognitive decline, yet the underlying damaging mechanisms remain incompletely understood. Here we established a positive correlation between plasma levels of serine proteinase elastase ELANE and periventricular white matter hyperintensity (PV-WMH) in a cohort of CSVD patients. In a CSVD murine model induced by bilateral carotid artery stenosis (BCAS), upregulated ELANE was detected both in microglia and peripheral blood neutrophils. Genetic ELANE deficiency significantly alleviated oligodendrocyte loss, thereby reducing white matter lesions (WMLs) as well as ameliorating sensorimotor and cognitive impairments in BCAS mice*. In vitro* studies demonstrated that ELANE triggered time-dependent and dose-dependent oligodendrocyte lineage cell death. Bone marrow transplantation showed that ELANE from microglia and peripheral blood both contributed to WML development and BCAS-induced neurological deficits. Mechanistically, ELANE, accumulated by oligodendrocytes, cleaved the phosphodiesterase domain of 2′,3′-cyclic nucleotide 3′-phosphodiesterase (CNPase). Pharmacological inhibition of ELANE with Sivelestat reduced oligodendrocyte loss and WMLs leading to the restoration of white matter integrity and neurological improvements in BCAS mice. In post-mortem brain specimens of CSVD patients ELANE accumulated within WMLs being predominantly localized in microglia (and hence defined as microglial ELANE) rather than in the brain-infiltrating neutrophils. We therefore posit microglial ELANE as an instigator of whiter matter injury in CSVD and suggest its potential therapeutic relevance.

## INTRODUCTION

Cerebral small vessel disease (CSVD) is widely present in elderly individuals accounting for 20%–25% of stroke as well as contributing to 45% of dementia cases worldwide ^[1–3]^. White matter lesions (WMLs), the most common feature of CSVD, are characterized by myelin degeneration and oligodendrocytes loss ^[4–6]^. Although much is known about the pathological features and clinical presentations of WMLs, the underlying cellular and molecular mechanisms are yet to be fully elucidated.

Oligodendrocytes, by myelinating axons in the central nervous system, are crucial for saltatory conduction of action potentials ^[7]^. In the adult brain, oligodendrocyte precursor cells (OPCs) maintain oligodendrocyte population at equilibrium and facilitate long-term repair of the white matter after injury or disease ^[8]^. Upon receiving demyelination signals, OPCs proliferate, migrate, populate the damaged regions, and differentiate into mature oligodendrocytes to restore myelin sheaths. OPC proliferation has been observed in ischemic WMLs after hypoperfusion injury ^[9]^; however, high susceptibility of OPCs and oligodendrocytes to cytotoxic and excitotoxic agents leads to their demise ^[10]^. Understanding the mechanisms responsible for the death of oligodendrocyte lineage cells is crucial for preventing myelin loss and arresting axonal degeneration in cases of white matter injury.

Neutrophils contribute to the white matter injury in the context of CSVD ^[11, 12]^. The neutrophil elastase ELANE, encoded by the *ELANE* gene, is a serine hydrolytic proteinase (EC 3.4.21.37) that is primarily synthesized in promyelocytes and stored in the cytoplasmic azurophil granules of neutrophils ^[13]^. Upon activation of neutrophils by external stimuli or during inflammation, ELANE is released into the extracellular matrix alongside other granule enzymes. ELANE is secreted by other cell types, such as monocytes, macrophages, platelets, smooth muscle cells, and microglia ^[14–16]^. The primary physiological role of ELANE is in host defense through degrading alien microorganisms or organic molecules phagocytosed by neutrophils ^[17]^. Although ELANE is beneficial for tissue turnover and infection control, its aberrant secretion in pathological conditions can mediate tissue damage in various diseases ^[18, 19]^. Intracerebral injection of ELANE disrupts the blood-brain barrier and triggers brain damage with intraparenchymal hemorrhage ^[20]^. Moreover, ELANE levels in plasma or cerebrospinal fluid are increased in patients with acute ischemic stroke or traumatic brain injury ^[21–23]^. Animal studies further indicated that ELANE plays a role in inflammatory responses after acute ischemic stroke or traumatic brain injury and contributes to neuronal cell death ^[24–26]^. Nonetheless, the detailed characterization of ELANE pathological features in the context of chronic brain injuries, including impairments of white matter tracts in CSVD, is, as yet, missing.

In this study, we revealed the role of microglial ELANE in the *in vivo* pathogenesis of WMLs and damage to oligodendrocyte lineage in CSVD patients as well as in a murine model of CSVD induced by bilateral common carotid artery stenosis (BCAS) ^[27]^.

## RESULTS

### Plasma ELANE positively correlates to periventricular white matter hyperintensity in patients with CSVD

The correlation between ELANE levels and clinical outcomes of the CSVD was studied in a cohort of 233 individuals diagnosed with CSVD. The mean age of the patients was 60.4 ± 11.5 years, and plasma ELANE levels were assessed using ELISA. The median ELANE level in plasma was 17.07 ng/mL (interquartile range: 12.82–22.20 ng/mL). Stratifying into tertiles based on ELANE levels showed that patients with higher ELANE levels demonstrated elevated diastolic blood pressure compared to those with lower ELANE readouts (Table S1).

Moreover, Pearson correlation analysis revealed a statistically significant positive correlation (r=0.14, P=0.04) between ELANE concentrations in patient’s plasma and periventricular white matter hyperintensity (PV-WMH), the latter being a measure of tissue damage (Table 1). This suggests a plausible involvement of ELANE in the development of WMLs in patients with CSVD.

**Table 1.**
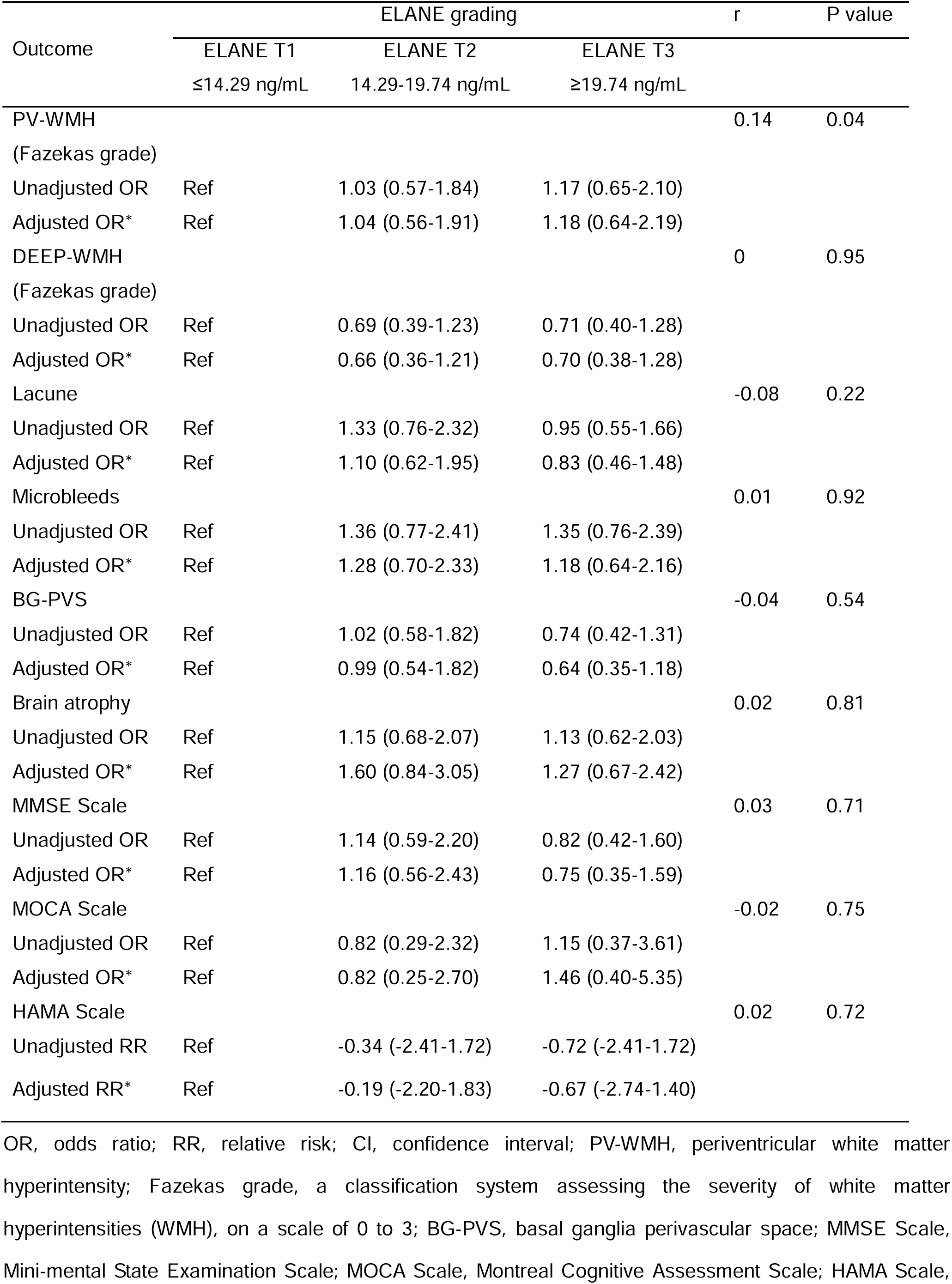
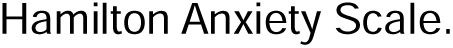
ORs or RR (95% CIs) for outcomes according to ELANE grading.

| Outcome | ELANE grading |  |  | r | P value |
| --- | --- | --- | --- | --- | --- |
|  | ELANE T1 | ELANE T2 | ELANE T3 |  |  |
|  | ≤14.29 ng/mL | 14.29-19.74 ng/mL | ≥19.74 ng/mL |  |  |
| PV-WMH<br>(Fazekas grade) |  |  |  | 0.14 | 0.04 |
| Unadjusted OR | Ref | 1.03 (0.57-1.84) | 1.17 (0.65-2.10) |  |  |
| Adjusted OR* | Ref | 1.04 (0.56-1.91) | 1.18 (0.64-2.19) |  |  |
| DEEP-WMH<br>(Fazekas grade) |  |  |  | 0 | 0.95 |
| Unadjusted OR | Ref | 0.69 (0.39-1.23) | 0.71 (0.40-1.28) |  |  |
| Adjusted OR* | Ref | 0.66 (0.36-1.21) | 0.70 (0.38-1.28) |  |  |
| Lacune |  |  |  | -0.08 | 0.22 |
| Unadjusted OR | Ref | 1.33 (0.76-2.32) | 0.95 (0.55-1.66) |  |  |
| Adjusted OR* | Ref | 1.10 (0.62-1.95) | 0.83 (0.46-1.48) |  |  |
| Microbleeds |  |  |  | 0.01 | 0.92 |
| Unadjusted OR | Ref | 1.36 (0.77-2.41) | 1.35 (0.76-2.39) |  |  |
| Adjusted OR* | Ref | 1.28 (0.70-2.33) | 1.18 (0.64-2.16) |  |  |
| BG-PVS |  |  |  | -0.04 | 0.54 |
| Unadjusted OR | Ref | 1.02 (0.58-1.82) | 0.74 (0.42-1.31) |  |  |
| Adjusted OR* | Ref | 0.99 (0.54-1.82) | 0.64 (0.35-1.18) |  |  |
| Brain atrophy |  |  |  | 0.02 | 0.81 |
| Unadjusted OR | Ref | 1.15 (0.68-2.07) | 1.13 (0.62-2.03) |  |  |
| Adjusted OR* | Ref | 1.60 (0.84-3.05) | 1.27 (0.67-2.42) |  |  |
| MMSE Scale |  |  |  | 0.03 | 0.71 |
| Unadjusted OR | Ref | 1.14 (0.59-2.20) | 0.82 (0.42-1.60) |  |  |
| Adjusted OR* | Ref | 1.16 (0.56-2.43) | 0.75 (0.35-1.59) |  |  |
| MOCA Scale |  |  |  | -0.02 | 0.75 |
| Unadjusted OR | Ref | 0.82 (0.29-2.32) | 1.15 (0.37-3.61) |  |  |
| Adjusted OR* | Ref | 0.82 (0.25-2.70) | 1.46 (0.40-5.35) |  |  |
| HAMA Scale |  |  |  | 0.02 | 0.72 |
| Unadjusted RR | Ref | -0.34 (-2.41-1.72) | -0.72 (-2.41-1.72) |  |  |
| Adjusted RR* | Ref | -0.19 (-2.20-1.83) | -0.67 (-2.74-1.40) |  |  |

OR, odds ratio; RR, relative risk; CI, confidence interval; PV-WMH, periventricular white matter hyperintensity; Fazekas grade, a classification system assessing the severity of white matter hyperintensities (WMH), on a scale of 0 to 3; BG-PVS, basal ganglia perivascular space; MMSE Scale, Mini-mental State Examination Scale; MOCA Scale, Montreal Cognitive Assessment Scale; HAMA Scale,
Hamilton Anxiety Scale.

### ELANE accumulates in the corpus callosum and plasma in BCAS mice

Three days after induction of BCAS, a murine model of CSVD, both concentration (Figure 1A) and activity (Figure 1B) of ELANE in the corpus callosum increased significantly compared to the sham group, and reached the peak at postoperative day 28. Flow cytometry analysis of tissues collected before BCAS showed that ELANE was expressed in mouse microglia and bone marrow cells (Figure S1). After BCAS, ELANE expression was elevated in microglia at postoperative day 7 (Figure 1C) and in bone marrow from postoperative day 3 (Figures 1D and 1E). Similarly to the corpus callosum, peak ELANE levels in bone marrow were observed at 28 days after BCAS induction coinciding with a decrease in brain myelin basic protein (MBP) (Figures 1D), the major constituent of oligodendrocyte myelin sheath, indicating progressive white matter injury. Furthermore, total ELANE concentrations in plasma increased continuously from 3 days after BCAS (Figure 1F). Thus, chronic WMLs resulting from CSVD are associated with an increased expression and activity of ELANE.

**Figure 1.**
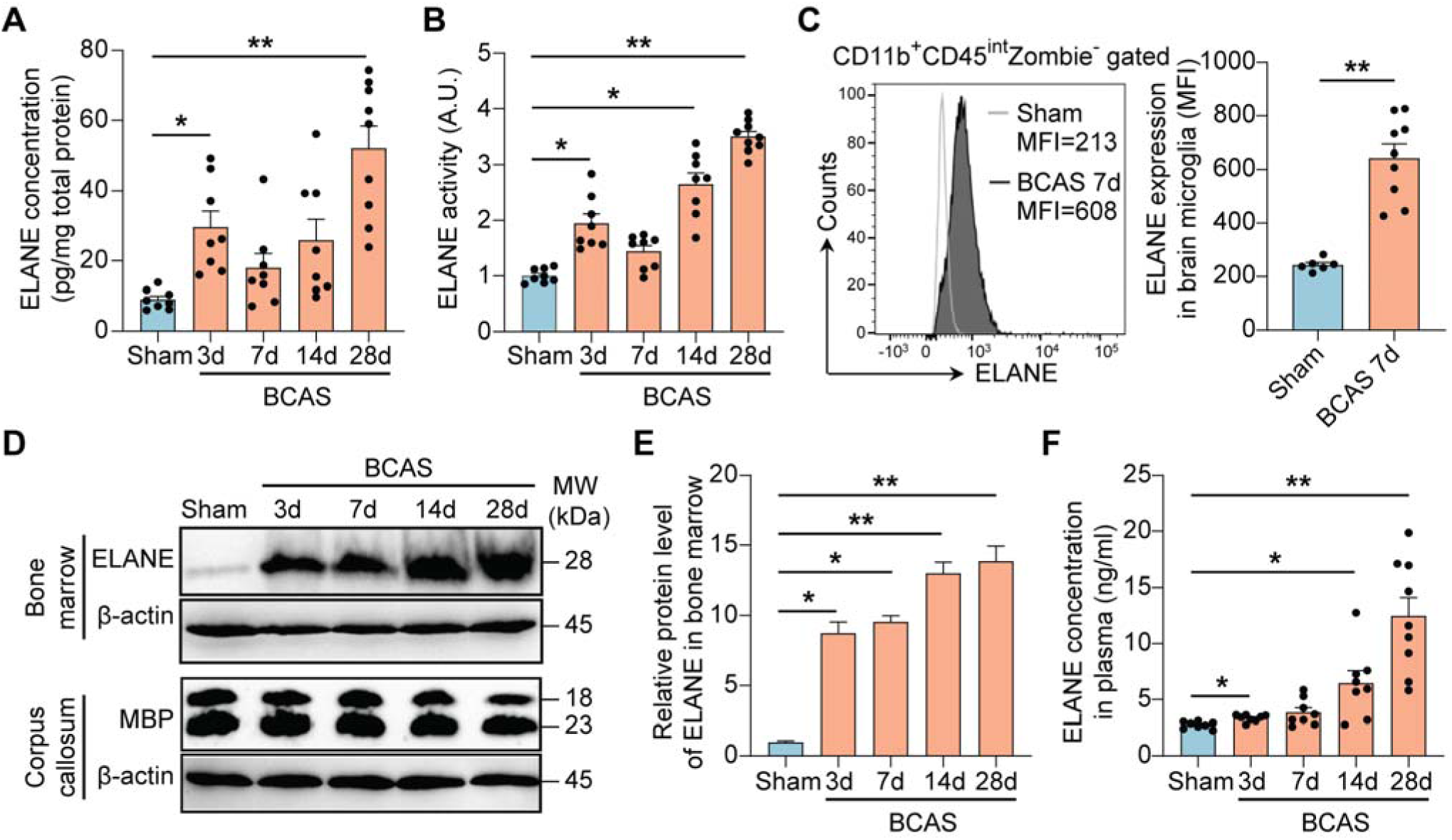
BCAS induces ELANE expression and myelin disruption in mice. (A) ELANE concentration in the mouse corpus callosum at different time points after BCAS surgery evaluated with ELISA. *n* = 8–9 mice per group. (B) ELANE catalytic activity in mouse corpus callosum homogenate samples at different time points after BCAS surgery measured by spectrophotometry. *n* = 8–9 mice per group. (C) ELANE expression in microglia was assessed by flow cytometric analysis at 7 days after BCAS or sham surgery. Quantifications of flow cytometry are displayed in the adjacent graph. *n* = 6–9 mice per group. (D) ELANE expression in the bone marrow, and MBP expression in the corpus callosum determined by western blot at 3 days, 7 days, 14 days, and 28 days after BCAS or sham surgery. β-actin served as an internal loading control. (E) Quantification of the intensity of ELANE expression in the bone marrow performed with ImageJ. *n* = 3 replicates per group. (F) Plasma ELANE concentration at different time points after BCAS or sham surgery evaluated with ELISA. *n* = 8–9 mice per group. See also Figure S1.

### *Elane* deletion protects against BCAS-induced white matter damage

We generated *Elane^−^*^/−^ (KO) mice (Figures 2A and 2B) and induced CSVD by BCAS. Cerebral blood flow after BCAS was similar in *Elane^−^*^/−^ mice and wild-type (WT) mice (P>0.05), suggesting that ELANE deficiency does not contribute to cerebral blood dynamics (Figure S2). At 28 days after BCAS, a notable reduction in MBP immunoreactivity was detected in the corpus callosum of WT mice, but not in *Elane^−^*^/−^ mice (Figure 2C). Staining with a highly sensitive indicator of myelin loss Luxol fast blue (LFB), revealed significant myelin impairment in the corpus callosum of WT BCAS mice compared to sham-operated controls; to the contrary myelin integrity was preserved in *Elane^−^*^/−^ BCAS mice (Figure 2D).

**Figure 2.**
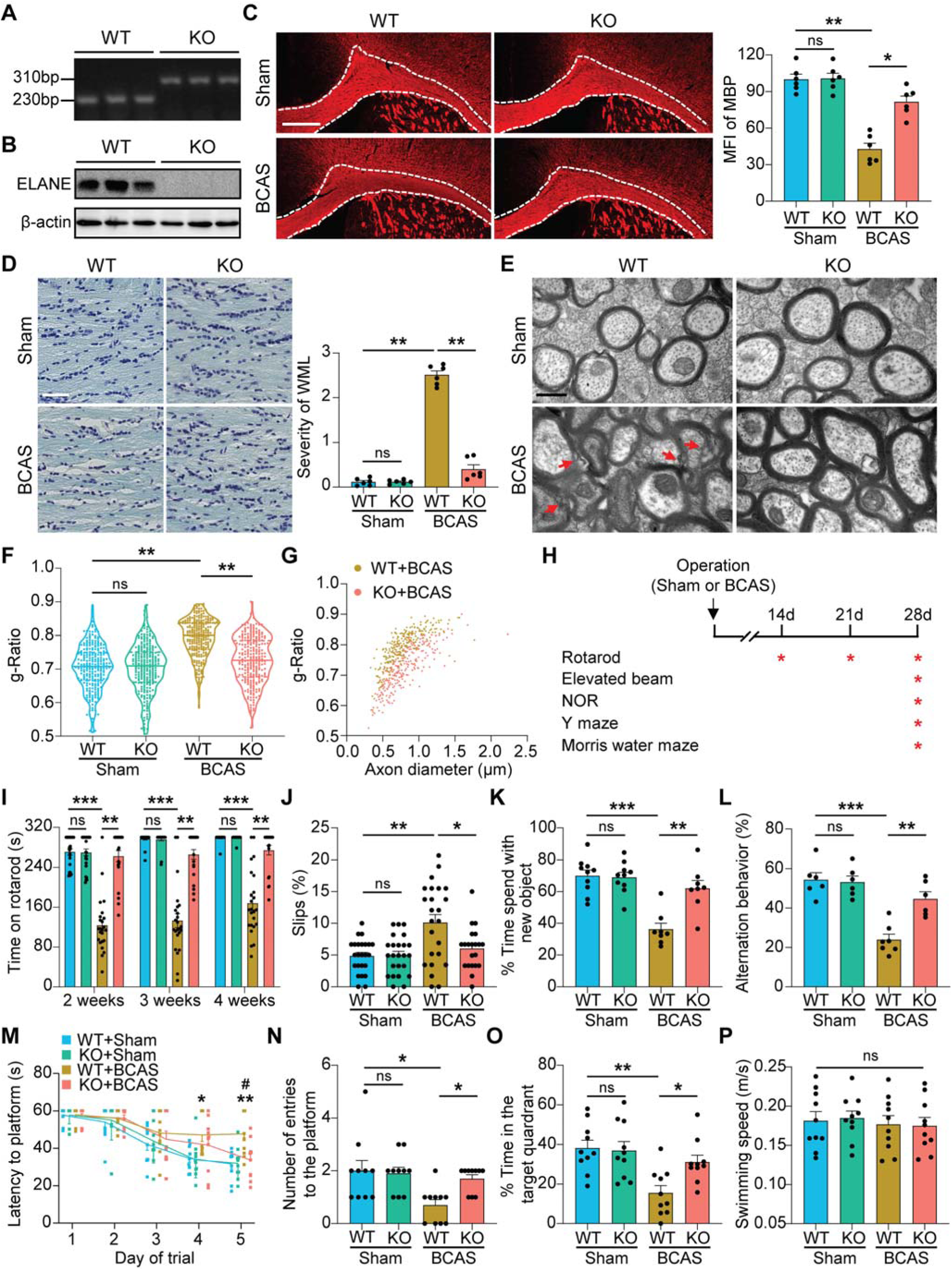
ELANE deficiency prevents corpus callosum white matter injury and protects against neurological impairment after BCAS surgery *in vivo*. (A) DNA extracts from tail tissue were subjected to PCR using primers flanking the loxp site and primers detecting *Elane* cDNA. (B) Expression of ELANE in mouse bone marrow was detected by western blot. β-actin served as an internal loading control. (C) Representative images of immunostaining for MBP and quantitative analysis of the mean fluorescence intensity of MBP in the corpus callosum. Scale bar, 400 μm. *n* = 6 mice per group. (D) Representative images of Luxol fast blue (LFB) staining and quantification of the severity of WMLs in the corpus callosum from WT and *Elane^−^*^/*−*^ mice at postoperative day 28. Scale bar, 50 μm. *n* = 6 mice per group. (E–G) Ultrastructural analyses of myelin integrity in the corpus callosum of indicated groups at postoperative day 28 using TEM. *n* = 3–4 mice per group. Axons used to calculate G-ratio: *n* = 250. (E) Representative electron microscopy images of WT and *Elane^−^*^/*−*^ mice on postoperative day 28. Red arrows indicate defective myelin sheaths. Scale bar, 1 μm. (F) Quantification of the G-ratios of myelinated axons. (G) Scatter plots of G-ratio as a function of axon diameter. (H) Experimental design for Figures 2I–2P. WT and *Elane^−^*^/*−*^ mice were subjected to neurological tests after BCAS or sham surgery. (I) Quantification of the latency to fall or spin around the rungs during the three trials for each mouse during the rotarod test. *n* = 7–8 mice per group. (J) Percentages of hind paw slipping for each mouse’s three runs on an elevated horizontal beam during the elevated beam test. *n* = 7–8 mice per group. (K) Quantification of the percentage of time spent exploring the novel object during the novel object recognition test session. *n* = 8–10 mice per group. (L) Quantification of the ratio of spontaneous alteration of each mouse in the Y-maze test. *n* = 6–7 mice per group. (M–P) The Morris water maze test. *n* = 10 mice per group. The latency to reach the platform (M) during the 5-day spatial learning test was recorded. In the memory probe test, the number of platform location crossings (N), the percentage of time spent in the platform quadrant (O), and swimming speed (P) were quantified. See also Figure S2.

Transmission electron microscopy (TEM) was employed to assess the ultrastructural integrity of myelin and axonal myelination in the corpus callosum of WT and *Elane^−^*^/−^ mice subjected to BCAS. Vacuoles and delamination were evident in WT mice 28 days after BCAS induction, whereas well-defined lamellar myelin sheaths remained intact in both sham-operated WT and *Elane^−^*^/−^ mice (Figure 2E). Myelin ultrastructure was also assessed by determining the G-ratio (axonal diameter/myelinated fiber diameter) on TEM images. The G-ratio was significantly increased in WT mice after BCAS compared to sham-operated controls (Figure 2F). In the *Elane^−^*^/−^ BCAS mice however, the G-ratio was not affected by BCAS (Figure 2F); moreover, the G-ratio remained low in *Elane^−^*^/−^ BCAS mice even when accounting for axon diameter (Figure 2G), highlighting a potential role of ELANE in the loss of myelination after BCAS. Thus, *Elane* deletion protects myelin sheath in the corpus callosum after BCAS.

### Deletion of ELANE prevents BCAS-induced neurological deficits

Given the essential role of white matter in preserving cognitive and motor function, we further examined the potential impact of ELANE on BCAS-induced motor and cognitive defects (Figure 2H). WT BCAS mice displayed a deficit in the accelerated rotarod test from 2 weeks after BCAS surgery compared to sham-operated controls, indicative of neurodegenerative behavioral patterns (Figure 2I). In contrast, *Elane^−^*^/−^ BCAS mice showed significantly greater time on the rotarod compared to WT BCAS mice (Figure 2I). Additionally, WT BCAS mice had impaired performance on a narrow-elevated beam test, which was preserved in *Elane^−^*^/−^ BCAS mice (Figure 2J).

We assessed cognitive performance of *Elane^−^*^/−^ BCAS and WT BCAS mice by novel object recognition, Y-maze, and Morris water maze tests. The novel object recognition test revealed an increase in exploratory behavior toward novel objects in *Elane^−^*^/−^ BCAS mice when compared to WT BCAS mice (Figure 2K). In the Y-maze test, WT BCAS mice had reduced alternative behavior, which was less pronounced in BCAS-operated *Elane^−^*^/−^ mice (Figure 2L). Morris water maze revealed preserved spatial learning and memory in *Elane^−^*^/−^ BCAS mice compared to WT BCAS mice at postoperative day 28, as evidenced by improvements in latency to the platform (Figure 2M), number of platform entries (Figure 2N), and time spent in the target quadrant (Figure 2O). No significant differences in swimming speed were observed among the groups (Figure 2P). Thus, ELANE deletion protected motor coordination and cognitive deficits induced by BCAS.

### ELANE deficiency prevents oligodendrocyte loss after BCAS

Previous research and our study support the notion that chronic hypoperfusion can lead to degeneration of oligodendrocytes linked to impaired myelin integrity (Figures 3A, 3B, and S3A) ^[28, 29]^. We thus quantified Olig2^+^ oligodendrocyte lineage cells in the corpus callosum. Our results showed that, at 28 days after BCAS, Olig2^+^ oligodendrocytes were greatly reduced in WT BCAS mice compared to sham controls, whereas they were preserved in *Elane^−^*^/−^ BCAS mice (Figures 3A, 3B, and S3A). Corpus callosum sections were co-stained for Olig2 and adenomatous polyposis coli clone CC1 (APC/CC1, a known marker of mature oligodendrocytes). At postoperative day 28, the number of Olig2^+^CC1^+^ cells in this region decreased in WT BCAS mice, which coincided with a decline in Olig2^+^ cell density, whereas cell numbers were preserved in *Elane^−^*^/−^ BCAS mice (Figures 3C, S3B, and S3C). Subsequent Olig2/NG2 co-staining was conducted to assess the impact of ELANE on OPCs. A notable decrease in Olig2^+^NG2^+^cells in the corpus callosum was observed in WT BCAS mice, but not in *Elane^−^*^/−^ BCAS mice at 28 days after BCAS surgery (Figures 3D, 3E, and S3D). These findings support direct involvement of ELANE in the loss of oligodendroglial lineage cells, which is a critical factor in white matter injury associated with CSVD.

**Figure 3.**
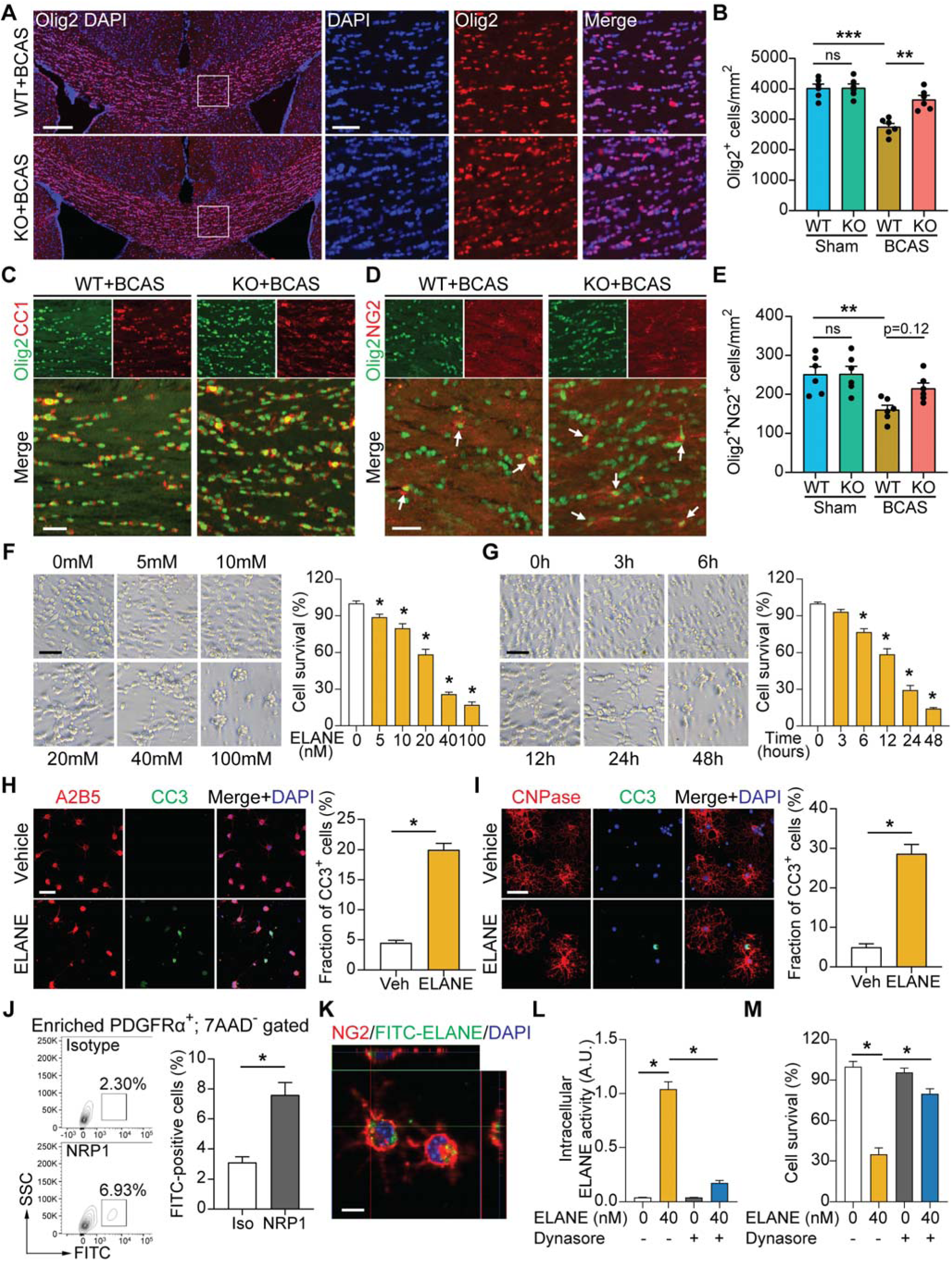
ELANE induces oligodendrocyte lineage cell loss *in vivo* and *in vitro*. (A) Representative images of white matter in the corpus callosum of WT and *Elane^−^*^/*−*^ mice that underwent BCAS surgery by immunostaining for Olig2 and with DAPI (the far left). Enlarged images of the boxed areas are on the right. Scale bars, 200 μm (left) and 20 μm (right). (B) Quantification of Olig2^+^ cells in the corpus callosum of WT and *Elane^−^*^/*−*^ mice that underwent BCAS or sham surgery. *n* = 6 mice per group. (B) Representative confocal images of co-immunostaining with Olig2 and CC1 antibodies in the corpus callosum. Scale bars, 50 μm. (C) Representative confocal images of corpus callosum labeled with Olig2 and NG2 antibodies. White arrows indicate double-stained cells. Scale bars, 50 μm. (D) Quantification of Olig2^+^NG2^+^ cells in the corpus callosum of WT and *Elane^−^*^/*−*^ mice that underwent BCAS or sham surgery. *n* = 6 mice per group. (E) Mouse OPCs were treated with ELANE at various concentrations for 24 hours. Representative light micrographs and quantification of viable cells using the CellTiter-Glo assay are shown. *n* = 6 replicates per group. Scale bar, 50 μm. (F) Mouse OPCs were treated with ELANE (40 nM) at various time points. Representative light micrographs and quantification of viable cells using the CellTiter-Glo assay are shown. *n* = 6 replicates per group. Scale bar, 50 μm. (G) Mouse OPCs were treated as indicated for 5 hours. Representative confocal images and quantification of CC3^+^ cells are displayed in the adjacent graph. *n* = 3 replicates per group. CC3, cleaved caspase-3. Scale bar, 50 μm. (H) Representative confocal images and quantification of CC3^+^ cells in differentiated mature oligodendrocyte culture treated with ELANE. *n* = 3 replicates per group. Scale bar, 50 μm. (I) Expression of NRP1 in mouse OPCs was assessed by flow cytometry. Representative images and quantification of NRP1 levels in PDGFRα-enriched OPCs are shown. *n* = 4 mice per group. (J) Representative confocal images of fluorescein-labeled ELANE (green) uptake by NG2^+^ OPCs (red). Nuclear staining with DAPI is shown in blue. Scale bar, 20 μm. (K) Intracellular ELANE activity of OPCs pre-treated with or without 60 μM Dynasore (30 minutes) was measured by spectrophotometry. *n* = 6 replicates per group. (L) Effect of ELANE on viability of OPCs pre-treated with or without 60 μM Dynasore (30 minutes). *n* = 6 replicates per group See also Figures S3 and S4.

### ELANE induces apoptosis of OPCs and mature oligodendrocytes *in vitro*

Next, we evaluated the effect of ELANE on oligodendrocytes viability by isolating and culturing primary murine OPCs *in vitro* (Figure S4A). Cell type verification was conducted using immunofluorescence staining with progenitor-specific antibodies to detect A2B5 and NG2 (Figure S4B). To assess the impact of ELANE on OPCs survival, primary murine OPCs were cultured in the presence of ELANE or vehicle. The number of OPCs significantly reduced in ELANE-treated cultures in a dose-dependent (Figure 3F) and a time-dependent (Figure 3G) manner compared to vehicle-treated controls. Furthermore, the proportion of cleaved caspase-3 (CC3)-positive A2B5^+^ cells was notably higher in ELANE-treated cultures compared to the vehicle-treated (Figure 3H), indicating that ELANE induces apoptosis. Then, mature differentiated oligodendrocytes derived from primary murine OPCs were identified by staining for MBP and CNPase (Figure S4C). We observed a concentration-dependent reduction in the population of mature oligodendrocytes after treatment with ELANE at 10 nM, 20 nM, 40 nM, and 100 nM, compared to untreated controls (Figure S4D). Moreover, immunostaining indicated a notable increase in the number of CC3^+^/CNPase^+^ apoptotic mature oligodendrocytes treated with ELANE compared to the control group (Figure 3I). Thus, ELANE induces apoptosis of OPCs and mature oligodendrocytes.

Previous studies showed that ELANE influences biological pathways through proteolysis ^[30]^. In the present study, we used phenylmethylsulfonyl fluoride (PMSF) to inactivate ELANE, and we confirmed that the treatment robustly decreased its catalytic activity (Figure S4E). Catalytically inactive ELANE did not reduce the number of OPCs *in vitro*, suggesting that ELANE-induced oligodendrocyte death is contingent with its catalytic activity (Figure S4F). The neuropilin 1 (NRP1), an ELANE receptor mediating ELANE uptake by cancer cells ^[31, 32]^, is also found in primary murine OPCs (Figure 3J); we also observed that ELANE was internalized and predominantly localized in the cytosol (Figure 3K). Treatment with Dynasore, a broad endocytosis inhibitor, reduced intracellular ELANE levels (Figure 3L) and protected OPCs against ELANE-induced cytotoxicity (Figure 3M). Thus, internalization of ELANE induces apoptosis of OPCs and mature oligodendrocytes.

### ELANE triggers apoptosis of OPCs and mature oligodendrocytes through cleavage of CNPase

CNPase (encoded by the *CNP* gene), a member of the 2H phosphodiesterase family, comprises approximately 4% of the total myelin protein in the central nervous system and is recognized as a structural protein predominantly present in OPCs and mature oligodendrocytes ^[33, 34]^. We repeated the aforementioned *in vitro* assays to determine the effect of supplementing the culture media with ELANE on CNPase degradation in oligodendrocytes and the impact of CNPase on ELANE-induced death of oligodendroglial cells. We observed a significant reduction in CNPase levels in both OPCs and oligodendrocytes in ELANE-treated compared to vehicle-treated cells (Figures 4A and 4B). Similarly, in a widely used oligodendrocyte precursor cell line CG4, ELANE induced a dose-dependent decrease in CNPase (Figure 4C). Consistent with the *in vitro* findings, CNPase protein levels were decreased in the corpus callosum of WT BCAS mice but not in *Elane^−^*^/−^ BCAS mice at postoperative day 28 (Figure 4D).

**Figure 4.**
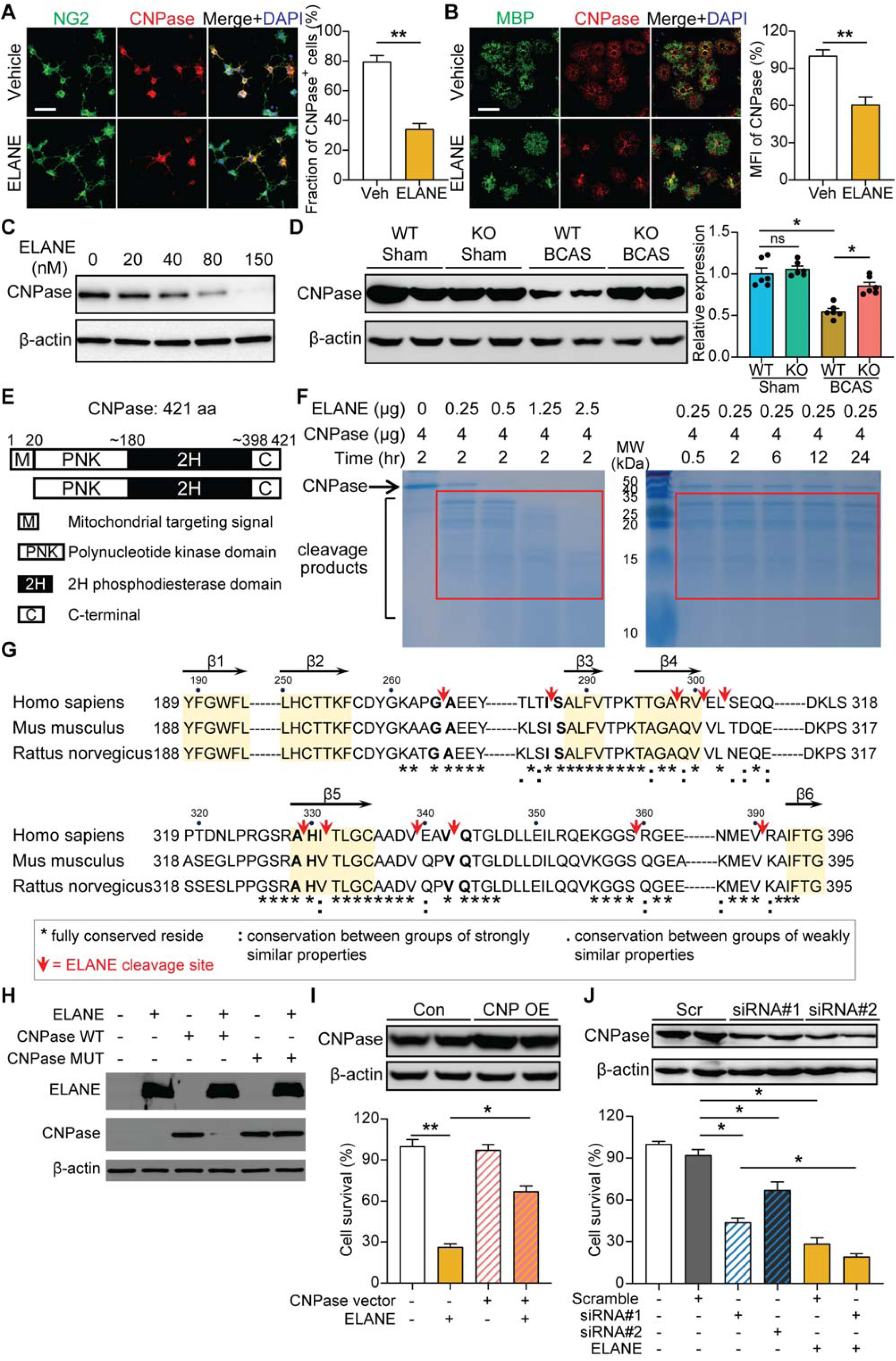
ELANE cleaves CNPase at the phosphodiesterase domain. (A) OPCs were treated as indicated for 5 hours and immunostained for NG2 (green) and CNPase (red). Nuclear staining with DAPI is shown in blue. Representative confocal images and quantification of the ratio of NG2^+^CNPase^+^ cells among total NG2^+^ cells are shown. *n* = 5 replicates per group. Scale bar, 50 μm. (B) Differentiated oligodendrocytes were treated as indicated for 5 hours. Representative confocal images labeled with MBP and CNPase are shown, and fluorescence intensity of CNPase staining is quantified. Data are calculated as fold change relative to the vehicle-treated group. *n* = 5 replicates per group. Scale bar, 100 μm. (C) Western blot to detect CNPase in CG4 cells incubated with the indicated concentrations of ELANE for 12 hours. (D) Expression of CNPase in the corpus callosum from WT and *Elane^−^*^/*−*^ mice at 28 postoperative day was determined by western blot. β-actin served as an internal loading control. The intensity of CNPase was quantified with ImageJ and is displayed in the adjacent graph. *n* = 6 mice per group. (E) Domain structure of CNPase. CNPase consists of a polynucleotide kinase domain, phosphodiesterase domain, and a C-terminal extension. In addition, isoform II contains an N-terminal mitochondrial targeting sequence (MTS), which is removed after mitochondrial import. PNK, polynucleotide kinase. 2H, phosphodiesterase domain featuring two conserved histidine residues. (F) Cleavage of recombinant human CNPase protein by indicated amounts of ELANE (left) or 0.25 μg of ELANE at indicated times (right) was assessed by SDS-PAGE and Coomassie blue staining. (G) Schematic representation of ELANE cleavage sites in CNPase. Red arrows indicate the position of ELANE cleavage within the phosphodiesterase domain of CNPase. Secondary structural elements in human CNPase are labeled. Conserved amino acid residues in human, mouse, and rat are indicated in bold. (H) Western blot to detect CNPase in HEK293T cells expressing *in vitro*-translated WT CNPase or cleavage-site mutant (MUT), and with or without co-transfected vector expressing *ELANE*. *In vitro*-translated ELANE, uncleaved WT CNPase, and uncleaved MUT CNPase are shown as references in lane 2, lane 3, and lane 5, respectively. (I) Western blot to detect CNPase in CG4 cells transfected with pCDH expressing CNPase or vector (upper). β-actin served as an internal loading control. Quantification of viability of CG4 cells overexpressing CNPase treated with ELANE as indicated (lower). *n* = 6 replicates per group. (J) Western blot analysis of endogenous CNPase expression after si-CNPase treatment for 48 hours in CG4 cells (upper). β-actin served as an internal loading control. After *CNPase* silencing, viable cells were quantified (lower). *n* = 9 replicates per group. See also Figures S5 and S6.

To identify cleavage sites recognized by ELANE in CNPase, we incubated recombinant human CNPase corresponding to full length of protein (amino acid [aa] 1–421, Figure 4E) with increasing concentrations of ELANE for 2 hours and evaluated protein size by SDS-PAGE electrophoresis. ELANE effectively bound and cleaved CNPase, while higher concentrations of ELANE yielded a higher proportion of smaller peptide fragments (Figures S5 and 4F). To determine the preferred cleavage sites, we incubated CNPase with 0.25 µg ELANE for 2 hours and then analyzed the peptides by N-terminal sequencing and mass spectrometry. This approach revealed 11 primary cleavage sites in human CNPase, all of which were located in the conserved 2H phosphodiesterase domain (Figures 4G and S6) and aligned well with ELANE’s sequence specificity as documented in the MEROPS database (https://www.ebi.ac.uk/merops). Examination of the human, murine, and rat CNPase aa sequence revealed the presence of ELANE cleavage sites at similar positions (Figure 4G). To verify these as ELANE cleavage sites, we co-expressed WT human CNPase or a CNPase mutant in which all 11 cleavage sites were mutated along with human ELANE in HEK293T cells, which do not endogenously express *CNP* and *ELANE*. This demonstrated the inability of ELANE to degrade the mutant CNPase protein (Figure 4H). Overexpression of full-length human CNPase in CG4 cells conferred protection against ELANE-induced cytotoxicity, potentially by increasing substrate availability (Figure 4I). Conversely, RNA silencing of CNPase in CG4 cells increased cell death and enhanced susceptibility to ELANE (Figure 4J). Collectively, these findings suggest that CNPase contributes to oligodendrocyte death triggered by ELANE.

### Brain-derived and peripheral blood-derived ELANE both contribute to BCAS-induced white matter injury

ELANE was initially identified at high levels in neutrophil azurophil granules, where it plays a crucial role in bacterial killing during acute infections ^[13]^. Additionally, microglia were shown to produce ELANE upon stimulation with lipopolysaccharide, LPS ^[16]^. To investigate the source of ELANE contributing to WMLs after BCAS, chimeric mice were generated to selectively express ELANE either in blood neutrophil or in microglia. Mononuclear cells (MNCs) from WT donor mice were transplanted into lethally myeloid depleted by irradiation *Elane^−^*^/−^ recipient mice to create chimeras expressing ELANE exclusively in peripheral blood (blood ELANE) (Figure 5A). Conversely, chimeric mice expressing ELANE solely in microglia (brain ELANE) were obtained by transplanting *Elane^−^*^/−^ MNCs into WT recipient mice. After verifying an average efficiency of more than 90% host blood cell replacement with donor cells (Figure 5B), these chimeric mice were subjected to BCAS. Demyelination was assessed 28 days after BCAS by MBP and LFB staining, which showed decreased MBP levels and increased myelin loss in the corpus callosum of both blood ELANE and brain ELANE chimeras compared to knockout (KO) chimeras (Figures 5C–5E). Similar reductions in the number of Olig2^+^ oligodendrocytes were also observed in the blood ELANE and brain ELANE chimeras compared to controls (Figures 5C and 5F). Furthermore, the average G-ratio of axonal fibers in blood ELANE and brain ELANE chimeras was higher than in KO chimeras, indicating increased myelin damage in both groups (Figures 5G and 5H). At day 28 after BCAS, motor balance performance significantly decreased in both blood ELANE and brain ELANE chimeras compared to KO chimeras (Figure 5I). Additionally, both blood ELANE and brain ELANE chimeras exhibited significantly reduced performance in alternation behavior compared to KO chimeras in the Y-maze test (Figure 5J). Thus, either brain- or blood-derived ELANE is sufficient to trigger WMLs and impaired neurological function after BCAS, implying a non-redundant role of brain-derived and blood-derived ELANE in the pathogenesis of white matter injury and neurological deficits in CSVD.

**Figure 5.**
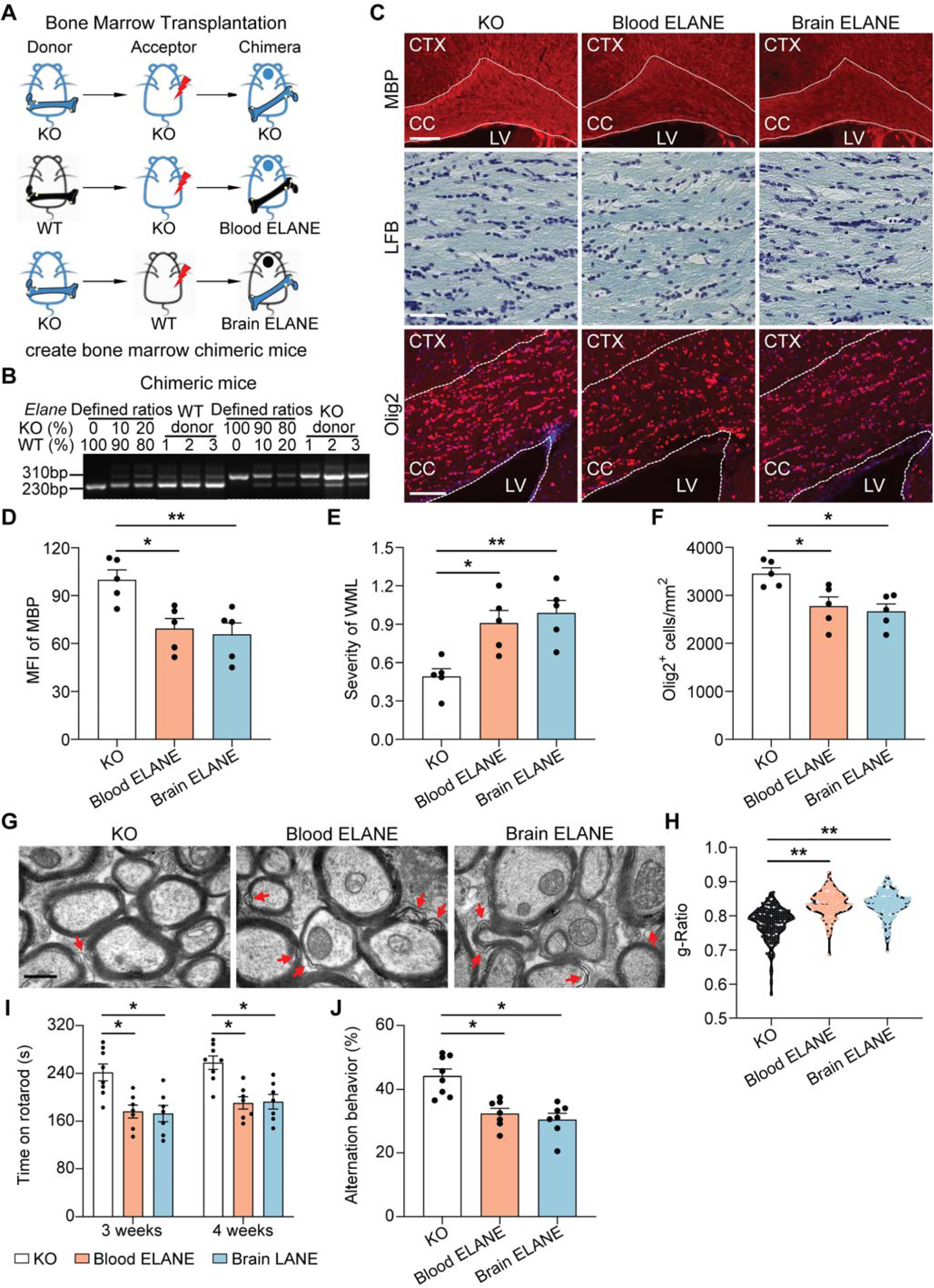
Brain-derived and bone marrow-derived ELANE both contribute to white matter injury after BCAS. (A) Schematic for generating blood ELANE or brain ELANE chimeric mice. Chimeric mice were generated to selectively express ELANE either in brain or peripheral blood and then underwent BCAS surgery. (B) Engraftment efficiency of chimeric mice was quantified using defined mixtures of WT and KO bone marrow cells. (C) Brain sections from BCAS-operated chimeric mice were labeled with MBP (upper), LFB (middle), and Olig2 (lower). The dotted white lines trace the boundaries of the corpus callosum. Scale bars, 200 μm (upper), 50 μm (middle), and 100 μm (lower). CC, corpus callosum; CTX, cortex; LV, lateral ventricle. (D) Quantification of mean fluorescence intensity of MBP in the corpus callosum are shown. MFI, mean fluorescence intensity. *n* = 5 mice per group. (E) The severity of WMLs was assessed by Luxol fast blue (LFB) staining. *n* = 5 mice per group. (F) Quantification of Olig2 in the corpus callosum on chimeric mouse brain sections. *n* = 5 mice per group. (G–H) TEM ultrastructural analysis of myelin integrity in the corpus callosum of chimeric mice at postoperative day 28. *n* = 3 mice per group. Axons used to calculate G-ratio: *n* = 200 (at least 50 axons per mouse). Representative TEM images (G) and quantification of the G-ratios of myelinated axons (H) in indicated groups. Red arrows indicate defective myelin sheaths. Scar bar, 1 μm. (I) Rotarod test was performed in BCAS-operated chimeric mice at indicated time points. The latency to fall or spin around the rungs for each mouse was quantified. *n* = 7–8 mice per group. (J) The ratio of spontaneous alteration of each mouse in the Y-maze test was quantified. *n* = 7–8 mice per group.

### Pharmacological inhibition of ELANE attenuates BCAS-induced white matter impairment and behavioral deficits

Sivelestat, a synthetic low-molecular-weight inhibitor of ELANE ^[35]^, was investigated for its therapeutic potential in inhibiting WMLs in BCAS mice. Treatment with Sivelestat resulted in a dose-dependent increase in MBP level, preservation of white matter integrity, and increased number of Olig2^+^ oligodendrocytes in the corpus callosum (Figures 6A–6D). Ultrastructural analysis revealed vacuoles and delamination in the corpus callosum of vehicle-treated BCAS mice 28 days after surgery, whereas mice treated with 10 mg/kg Sivelestat maintained intact lamellar myelin sheaths (Figure 6E). Sivelestat also protected myelin, as mice treated with 10 mg/kg Sivelestat had much lower G-ratios (Figures 6F and 6G), although the overall axon diameter did not change between the treatment groups (Figure 6H). Mice treated with Sivelestat after BCAS surgery exhibited a dose-dependent enhancement of motor coordination and cognitive function compared to the vehicle-treated group, with a significant improvement observed at a high dose of 10 mg/kg (Figures 6I–6K). These findings demonstrate that pharmacological inhibition of ELANE by Sivelestat may partially reverse sensorimotor deficits and spatial working memory impairments in BCAS-operated mice.

**Figure 6.**
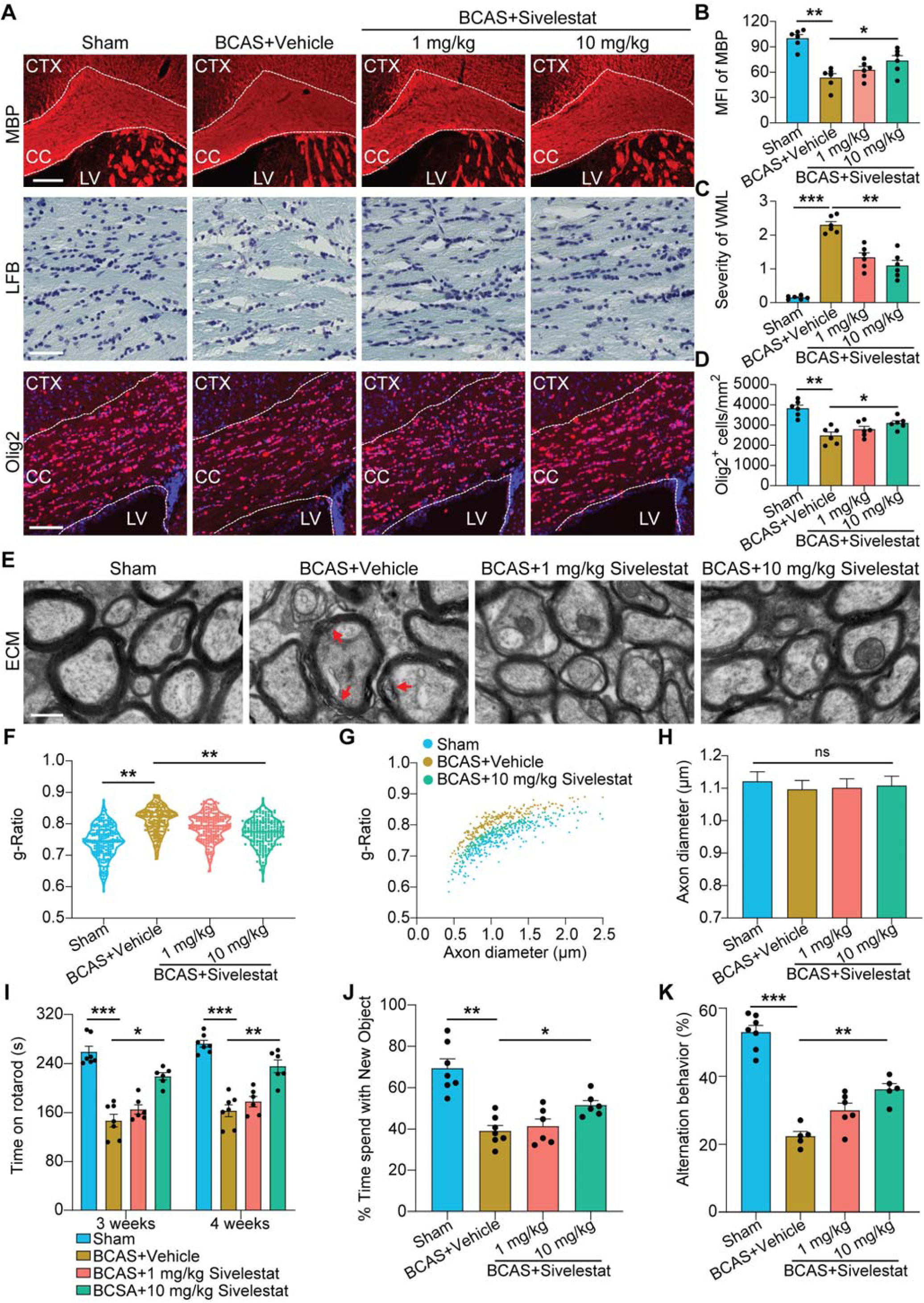
Administration of ELANE inhibitor Sivelestat attenuates white matter damage and neurodeficits in BCAS mice. (A) Representative images of coronal sections labeled with MBP (upper), LFB (middle), and Olig2 (lower) from vehicle or Sivelestat-treated BCAS- or sham-operated mice. Dotted white lines indicate the corpus callosum boundaries. Scale bars, 200 μm (upper), 50 μm (middle), and 100 μm (lower). CTX, cortex; CC, corpus callosum; LV, lateral ventricle. (B) Quantification of MBP mean fluorescence intensity in the corpus callosum. *n* = 6 mice per group. (C) Quantification of the severity of WMLs in the corpus callosum area. *n* = 6 mice per group. (D) Quantification of Olig2^+^ cells in the corpus callosum. *n* = 6 mice per group. (E) Representative electron microscopy images in the corpus callosum at postoperative day 28 in Sivelestat-treated and vehicle-treated BCAS mice. Red arrows indicate defective myelin sheaths. Scale bar, 1 μm. (F) Quantitative analysis of the G-ratios of myelinated axons. *n* = 200 myelinated axons (at least 50 axons per mouse, three mice per group) for each group. (G) Scatter plots of G-ratio as a function of axon diameter. *n* = 200 axons per group (at least 50 axons per mouse, three mice per group). (H) Quantification of axonal diameter in indicated groups. *n* = 200 axons per group (at least 50 axons per mouse, three mice per group). (I) Quantification of rotarod test results at indicated time points. *n* = 6–7 mice per group. (J) Percentage of time spent exploring the novel object during the novel object recognition test session was quantified. *n* = 6–7 mice per group. (K) The ratio of spontaneous alteration of each mouse in Y-maze test was quantified. *n* = 5–7 mice per group.

### ELANE accumulates in the corpus callosum microglia in patients with CSVD

Finally, we examined the expression pattern of ELANE in postmortem brain tissues from CSVD patients and compared it with age- and sex-matched healthy individuals. The ELANE expression was markedly increased in the corpus callosum from CSVD patients compared to control individuals, where ELANE^+^ cells were scarce (Figure 7A). We employed a previously described classification scheme to categorize the ELANE^+^ cells; specifically, round cells with intracellular ELANE were classified as ‘inactive’ (Figure 7B, i), whereas cells exhibiting elongation indicative of cell migration (Figure 7B, ii) or degranulation with extracellular ELANE (Figure 7B, iii) were classified as ‘active’ ^[36]^. Remarkably, 87.9%±12.9% (*n* = 8) of ELANE^+^ cells in patients with CSVD exhibited an ‘active’ phenotype (Figure 7B), supporting the assertion that ELANE expression is increased in WMLs in patients with CSVD. Co-staining of ELANE indicated colocalization of ELANE^+^ cells with microglia and brain-infiltrating neutrophils in the corpus callosum of patients with CSVD (Figures 7C and 7D). Specifically, ELANE expression was most abundant in Iba1^+^ brain microglia rather than in neutrophils (Figure 7E). Therefore, ELANE builds up in the brains of patients with CSVD, is primarily associated with microglia. This suggests a significant involvement of microglial-derived ELANE in the white matter injury observed in CSVD patients.

**Figure 7.**
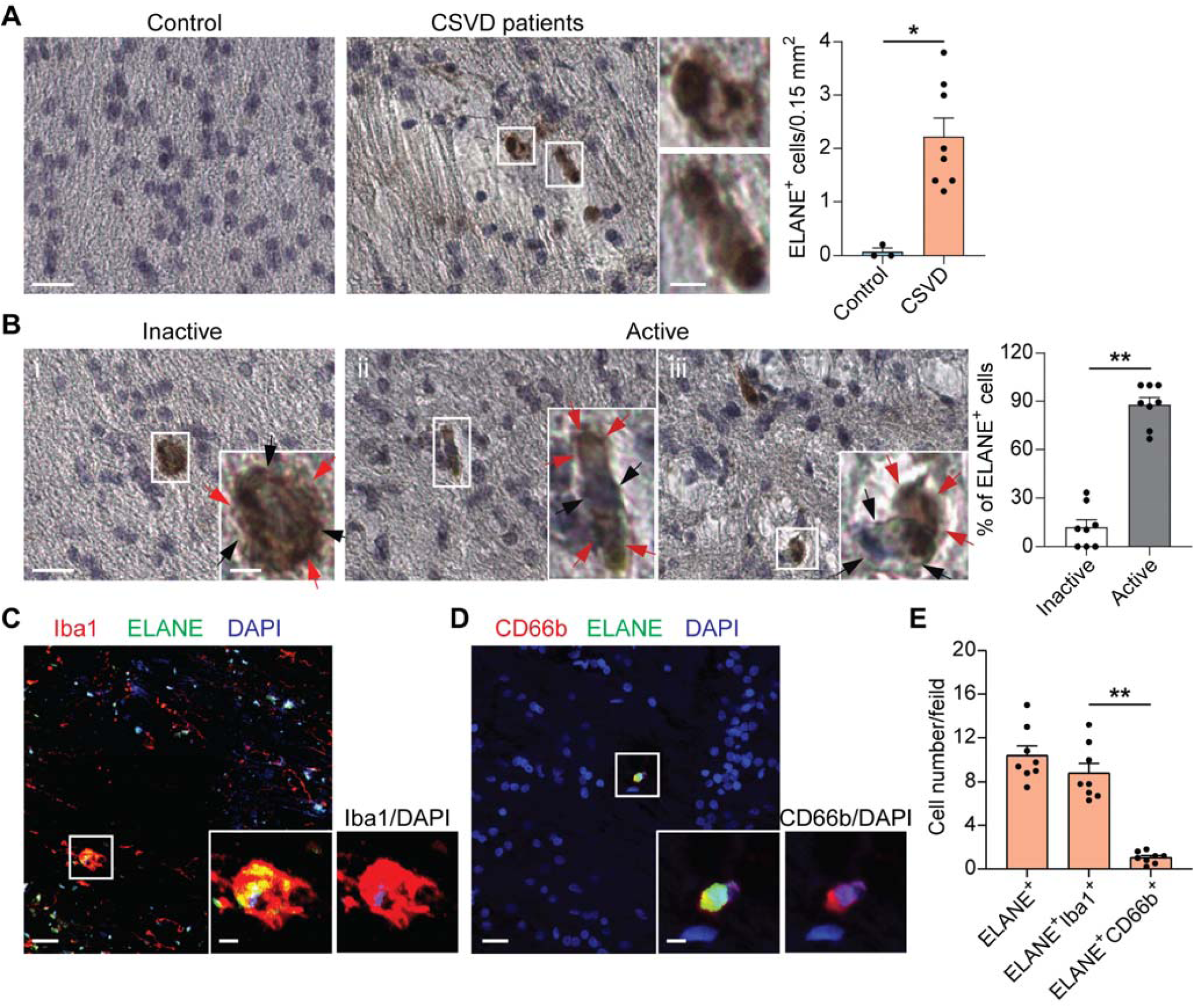
ELANE accumulates in the corpus callosum of CSVD patients. (A) Representative images of ELANE^+^ staining in the corpus callosum from control individuals or patients with severe CSVD. Images were taken at 40× magnification. *n* = 3–8 per group. White squares indicate the ELANE^+^ cells enlarged in the right panel images. Quantification of ELANE^+^ cells is displayed in the adjacent graph. Scale bar, 20 μm (left) and 5 μm (right). (B) Inert and active ELANE^+^ cells in the corpus callosum from patients with CSVD. (i) ELANE (red arrows) is intracellular (black arrows). (ii) Elongated ELANE^+^ cells (red arrows). (iii) Extracellular ELANE (red arrows). Quantification is displayed in the adjacent graph (*n* = 8 per group). Scale bar, 20 μm and 5 μm (insert). (C) Immunostaining of ELANE (green) in brain Iba1^+^ microglia (red) in corpus callosum from patients with CSVD. Scale bar, 20 μm and 5 μm (insert). (D) Immunostaining shows brain-infiltrating neutrophil expression of CD66b (red) and ELANE (green) in corpus callosum from patients with CSVD. Scale bar, 20 μm and 5 μm (insert). (E) Quantification of different ELANE^+^ cell types in brain sections from patients with CSVD in corpus callosum area (*n* = 8 per group).

## DISCUSSION

This study reveals the fundamental role of ELANE accumulation in the corpus callosum of CSVD patients in triggering white matter damage. Genetic or pharmacological suppression of ELANE substantially enhanced the survival of oligodendroglial lineage cells, mitigated the progression of white matter injury, and led to enduring improvements in behavioral outcomes after hypoperfusion injury. We further demonstrate that ELANE-induced oligodendrocyte death and demyelination involve proteolytic cleavage of CNPase at multiple sites. These results underscore the causal involvement of ELANE in driving oligodendrocyte lineage demise and WML pathology in the context of CSVD.

ELANE is known to be expressed in bone marrow precursor cells at the promyelocyte stage in peripheral circulation and in microglia in the central nervous system ^[16, 37–39]^. Disruption of the blood-brain barrier after hypoperfusion permits the infiltration of leukocytes, including neutrophils, into the white matter ^[12]^, implicating neutrophils as a potential source of elevated brain ELANE levels. Additionally, ELANE released by microglial cells can directly interact with extracellular matrix substrates and various brain cells, suggesting a similar role for microglia-derived ELANE during brain injury ^[39]^. We confirmed a significant linear relationship between plasma ELANE levels and periventricular white matter hyperintensity (a measure of white matter damage) in CSVD patients. Specifically, higher plasma ELANE levels correlate with higher Fazekas grades of PV-WMH, the latter indicating more severe damage to brain white matter. Our research further demonstrated that the accumulation of ELANE protein in the affected human brain tissue primarily originates from microglia. As WMLs progress, ELANE expression increases in both bone marrow and microglia, and ELANE derived from the brain induces white matter damage and loss of oligodendrocytes comparable with ELANE derived from bone marrow. Collectively, our study highlights the causal role of microglia-derived ELANE (microglial ELANE) in conjunction with peripheral neutrophil elastase in causing oligodendrocyte death and myelin destruction, ultimately resulting in WMLs. Further research is needed to differentiate the shared and distinct functions of neutrophil elastase and microglial elastase.

We also discovered that ELANE directly triggers cell death of OPCs and mature oligodendrocytes, leading to WMLs and subsequent functional impairment. In the adult brain, OPCs are recruited as part of an endogenous response to fill the damaged areas and differentiate into mature oligodendrocytes to restore myelin sheaths ^[40, 41]^. Our findings demonstrate that ELANE induces OPCs death in a dose- and time-dependent manner, and that inhibition of ELANE mitigates damage to OPCs and mature oligodendrocytes in the injured corpus callosum after BCAS surgery. The pro-apoptotic impact of ELANE is partially reversed by Dynasore, implicating endocytosis in ELANE-mediated oligodendrocyte proteolysis. In addition to its direct cytotoxic effects on oligodendrocytes, ELANE may compromise the blood-brain barrier, recruit immune cells, and provoke the release of pro-inflammatory cytokines, exacerbating secondary brain damage ^[20, 39]^. Additional studies are needed to elucidate the influence of these indirect factors on oligodendrocytes loss and WML development after CSVD.

Another discovery of this study is the characterization of ELANE cleavage sites on the CNPase protein, leading to its degradation. This finding may partially explain the observed decrease in CNPase levels and reduced myelination in the brains of patients with Alzheimer’s disease and multiple sclerosis ^[42–44]^, accompanied by elevated levels of ELANE ^[45, 46]^. We identified that the preferred ELANE cleavage sites are mainly in the conserved 2H phosphodiesterase domain of CNPase, which is necessary for its enzymatic phosphodiesterase activity ^[47]^; while 11 preferred sites were found in this domain, other biologically relevant cleavage sites may also exist. It was reported that CNPase binds to tubulin and that it plays a cytoskeletal role in anchoring microtubules to the plasma membrane as well as in regulating tubulin polymerization ^[48]^. CNPase is also known to catalyze the hydrolysis of toxic 2′,3′-cyclic nucleotides to neuroprotective 2′-adenosine monophosphate ^[44]^. Also, 2′,3′-cyclic nucleotides were found to promote opening of mitochondrial permeability transition pores, leading to apoptosis and necrosis ^[49–51]^. As expected, we found that silencing of *Cnpase* expression in rat oligodendrocyte progenitor CG4 cells caused cell death, and we also confirmed the dependence of ELANE-induced oligodendrocyte death on CNPase. Our findings may help to further define the critical role of CNPase in oligodendrocytes, where it is highly expressed.

We observed that the detrimental effects of ELANE on oligodendrocytes and white matter integrity after BCAS were partially mitigated by the highly specific small molecule inhibitor of ELANE, Sivelestat. Treatment with Sivelestat resulted in a dose-dependent reduction in WMLs while also preserving oligodendrocytes and improving behavioral outcomes. These findings are in line with the previous study demonstrating that Sivelestat mitigates myelin loss in neuromyelitis optica (NMO) by inhibiting neutrophil migration into the NMO lesion ^[36]^. Our research contributes to the understanding of how Sivelestat alleviates ELANE-induced damage to oligodendrocytes, a key factor in myelin loss. Sivelestat is currently approved for clinical use in Japan and South Korea for patients with acute lung injury associated with systemic inflammatory response ^[52, 53]^, and it is also approved by the China Food and Drug Administration for clinical use in coronavirus disease 2019-related acute respiratory distress syndrome. Being a cell-permeable small molecule, Sivelestat can inhibit intracellular ELANE in both circulating neutrophils and resident microglia ^[35, 54]^. These initial findings suggest that Sivelestat might be regarded as a potential treatment for white matter diseases linked to oligodendrocyte loss. However, further comprehensive preclinical studies are essential before considering the translation of this approach into clinical practice.

In summary, this study provides new insights into the pathogenetic mechanism of microglia-derived ELANE acting in conjunction with peripheral neutrophil elastase to induce oligodendrocyte damage and the development of WMLs, identifying ELANE as a promising therapeutic target for managing small vessel disease and vascular dementia.

## ACKNOWLEDGMENTS

This work was supported in part by National Natural Science Foundation of China (82001243, 82122021).

## AUTHOR CONTRIBUTIONS

Y.W., F.-D.S., and W.-N.J. formulated the study concept. C.-Y.D., W.C., Q.L., L.S., Y.W., M.S., Y.H., J.X., Y.C., N.W., Y.H., and L.J. performed the experiments. W.-N.J., C.-Y.D., Q.L., Y.P., J.F., W.D., T.L., and W.H. analyzed the data and interpreted the results. W.-N.J., C.-Y.D., and W.C. wrote the paper. Y.W., F.-D.S., A.V., K.S., X.M., and Y.W. discussed, edited, and contributed to the writing.

## DECLARATION OF INTERESTS

The authors declare no competing interests.

## METHODS

### Study participants

This study recruited 233 patients with CSVD from the Department of Neurology at Beijing Tiantan Hospital between 2019 and 2024. The study was approved by the Ethics Committee of Beijing Tiantan Hospital, Capital Medical University (Approval Number: KY201914002, Approval Date: December 29, 2019). This research was conducted in accordance with the World Medical Association’s Declaration of Helsinki, and all patients provided written informed consent before being enrolled in the study. The inclusion criteria were: were: (1) at least 18 years old; (2) sign the informed consent form; (3) have independence in daily life (with a mRS score ≤2; and (4) have visible SVD lesions on head MRI. For inclusion criterion (4), any of the following conditions could be met to qualify for the study: high signal in the white matter of the brain with a Fazekas score ≥2; a Fazekas score of 1 accompanied by two or more vascular risk factors (including but not limited to hypertension, hyperlipidemia, diabetes, obesity, current smoking, or a history of vascular events other than stroke); a Fazekas score of 1 with concurrent lacunar infarction; or imaging indicating new subcortical lacunar infarction.

The exclusion criteria were (1) acute cerebral infarction (lesions showing high signal on DWI with a diameter >20mm), (2) acute cerebral hemorrhage, (3) acute subarachnoid hemorrhage, (4) history of cerebrovascular malformation or aneurysmal subarachnoid hemorrhage, (5) presence of untreated aneurysms, (6) confirmed neurodegenerative diseases, (7) definite non-vascular white matter lesions, (8) psychiatric disorders diagnosed according to DSM-V criteria, (9) contraindications for MRI examination, (10) severe organic diseases with an expected survival time of less than 5 years, (11) inability to complete follow-up due to geographical or other reasons, and (12) concurrent participation in other clinical trials.

### Assessment of demographic and clinical information

Demographic and clinical information was obtained from medical charts and patients’ self-descriptions, including age, sex, body mass index (BMI), history of hypertension, diabetes, dyslipidemia, smoking habits, history of transient ischemic attack (TIA), acute single or multiple ischemic infarctions, modified Rankin Scale (mRS) score, and National Institutes of Health Stroke Scale (NIHSS) score.

### Brain MRI acquisition and evaluation

MRI scan (3.0 T) was conducted using the following sequences: T1-weighted imaging, T2-weighted imaging, FLAIR imaging, and either susceptibility-weighted imaging (SWI) or gradient-recalled echo (GRE) imaging. The neuroimaging biomarkers of CSVD were defined based on the STandards for ReportIng Vascular changes on nEuroimaging (STRIVE). The CSVD burden score was determined by assigning one point for each of the following MRI parameters: severe white matter hyperintensities (WMH) (periventricular WMH Fazekas score of 3 or deep WMH Fazekas score of 2–3), the presence of lacunes, microbleeds (MBs), and moderate-to-severe basal-ganglia perivascular spaces (BG-PVS, *N* > 10). The total score ranged from 0 to 4 points. For group comparisons, patients were further classified into those with no/mild CSVD burden (burden score 0–1) and those with moderate/severe CSVD burden (burden score 2–4). This stratification aligned with the distribution of CSVD profiles observed in our data and previous studies. All images were evaluated by two blinded readers who were unaware of the patient information.

### Cognitive assessments

All patients completed cognitive assessments using the Mini-Mental State Examination (MMSE), Montreal Cognitive Assessment (MoCA), Hamilton Anxiety Scale (HAMA) and Hamilton Depression Scale (HAMD).

### Postmortem human brain tissues

Collection of human samples was performed according to protocols approved by the institutional review board of Beijing Tiantan Hospital (Beijing, China). The pathologic diagnosis was based on pathologic diagnostic criteria for CSVD made by a neuropathologist. For immunostaining of human brain tissues, three cases of control postmortem and eight CSVD human brain sections were evaluated. We studied the corpus callosum area in all cases. Brain tissues for control cases were obtained from deceased individuals with non-neurologic diseases and without history of neurological or neuropsychiatric conditions. Tissue sections of 5-μm thickness were cut and immunostained with ELANE antibody (ab68672, Abcam, Cambridge, UK). To quantify the expression of ELANE in the brain, we examined five randomly selected microscopic fields in the corpus callosum of each brain slide and counted ELANE^+^ cells in a 0.15-mm^2^ field of the corpus callosum. ELANE^+^ cells were classed as “inactive” versus “active” based on their morphology as previously reported ^[36]^.The data were analyzed by researchers blinded to experimental conditions. The average age of CSVD and control subjects did not differ significantly (CSVD: 57.7 ± 2.19 years old [y/o]; control: 61.8 ± 3.42 y/o; means ± s.e.m.; *P* > 0.05; unpaired *t*-test]).

### Mice

All animal experiments were performed in accordance with ARRIVE (Animal Research: Reporting In Vivo Experiments) guidelines. All animal experiments were performed in accordance with protocols approved by the animal care and use committee of Beijing Tiantan Hospital. C57BL/6J and C57BL/6N mice were purchased from Beijing Vital River Laboratory Animal Technology Co., Ltd. (Beijing, China). *Elane^−/−^* mice (JAX: 006112) were purchased from The Jackson Laboratory (Bar Harbor, Maine). All mutant mice were backcrossed to the C57BL/6 background for at least 12 generations. Animals were housed under pathogen-free conditions, with a maximum of five animals per cage, standardized light/dark cycle conditions, and free access to food and water. Genotyping for *Elane^−/−^* mice was performed using the following primers (5′ - 3′): common-TGCACAGAGAAGGTCTGTCG, WT forward-GGAACTTCGTCATGTCAGCA; mutant forward-TGGATGTGGAATGTGTGCGAG.

### BCAS surgical procedure and drug administration

Male C57BL/6 mice (7–9 weeks old, 22–26 g) were anesthetized with 4.0% isoflurane and maintained on 1.5% isoflurane in 70% N_2_O and 30% O_2_ using a small animal anesthesia system. After a midline cervical incision, both common carotid arteries were exposed. A micro-coil with a diameter of 0.18 mm (Sawane Spring Co., Shizuoka, Japan) was implanted surgically on the bilateral common carotid artery. Rectal temperature was maintained between 36.5 °C and 37.5 °C.

In the Sivelestat efficacy study, four groups each consisting of 5–7 animals were used. Sivelestat (1 mg/kg and 10 mg/kg) (ONO-5046, catalog no. 3535, Tocris) was administered intraperitoneally into male C57BL/6 mice 24 hours before the BCAS surgical procedure and once daily for 4 weeks after the BCAS surgery. The vehicle group similarly received intraperitoneal injections of PBS. In the sham group, mice underwent a sham surgery.

### Bone marrow transplantation

To induce bone marrow chimeric mice, 8-week-old male recipient mice (C57BL/6J WT or *Elane^−/−^* mice) were exposed to a single lethal dose of 10 Gy ^60^Co irradiation 24 hours before the bone marrow transplant (BMT). A single-cell bone marrow suspension was prepared from hind leg tibias and femurs of donor *Elane^−/−^* or C57BL/6J WT mice. Then, 1 × 10^6^ cells were intravenously injected into recipient C57BL/6J WT or *Elane^−/−^* mice to generate brain ELANE or blood ELANE chimeric mice. Recipient mice received drinking water supplemented with fluoroquinolone antibiotic enrofloxacin (0.01% Baytril, Bayer Vital) for 5 weeks.

Polymerase chain reaction (PCR) detection of the *Elane^−/−^* genotype was performed on DNA extracted from blood of recipient mice 10 weeks after transplant to confirm the efficiency of this BMT technique. Bone marrow reconstitution of recipient mice was further confirmed by PCR analyses of *Elane* mRNA expression levels in bone marrow mononuclear cells at the end of the procedure.

### Cerebral blood flow measurement

Laser-speckle contrast imaging was used to measure cortical cerebral perfusion at baseline and 28 days after sham or BCAS surgery. Under anesthesia induced with 1.5% isoflurane in 70% N_2_O and 30% O_2_, a 2.0-mm-diameter laser Doppler flowmetry probe was directly placed on the skull 1.0 mm posterior to bregma and 2.5 mm lateral to midline. Body temperature was monitored throughout and maintained at 37 °C ± 0.5 °C using a heat pad. After measurement, the skin on the skull bone was sutured. These procedures were repeated for every period of cerebral blood flow (CBF) measurement. Baseline CBF was obtained immediately before the surgery, and CBF was measured 28 days after the surgery. CBF data for each animal were calculated as a percentage with respect to the baseline CBF.

### Morris water maze test

The Morris water maze test was performed in a circular pool (diameter = 100 cm) with a square platform (11 × 11 cm^2^) submerged 2 cm beneath the water surface. An edible pigment was used to create opacity of the water to camouflage the submerged platform. Spatial acquisition trials were conducted over five consecutive days. Mice were placed into the pool from one of four designated start positions and allowed to locate the hidden platform for 60 seconds. Each mouse was trained on four trials per day with an interval of 15 seconds between trials. At the end of each trial, the mouse was placed on the platform or allowed to stay on the platform for 30 seconds with prominent spatial cues displayed around the room. The latency was defined as the time that the animal spent to reach and climb onto the platform from the start position. On day 6, the platform was removed, and a single 60-second probe trial was conducted. The percentage of time spent in the goal quadrant and the number of times that the animal crossed the zone where the platform was previously located was recorded.

### Object recognition test

To evaluate non-spatial working memory, an object recognition test was performed as previously described ^[55]^. Mice were placed in a 30 cm (long) × 30 cm (wide) × 30 cm (height) box and were permitted to explore the testing area for 10 minutes per day over a course of 3 days. On the day of the test, a session with two trials was completed. The intertrial interval was 6 hours. In the first trial, two same objects were placed in the box, and mice were allowed to freely interact with the objects for 10 minutes. Exploration was considered as directing the nose at a distance <1 cm from the object and/or touching it with the nose. In the second trial, one of the objects was replaced with a new one, and the mice explored the testing area once again for 10 minutes. Discrimination time was computed as the time spent at the old object subtracted from the time spent at the novel one and was considered an indicator of recognition memory.

### Y-maze test

To evaluate spatial working memory, the Y-maze spontaneous alteration test was carried out using a plastic Y-shaped runway. The Y-shaped runway had three arms (40 cm long, 12 cm high, and 3 cm wide) diverging at a 120° angle from the central point. The arm where the mice were initially placed was labeled A, with the other two arms being labeled B and C. Experiments were performed over a 10-minute session, and the total number and sequence of arm entries were recorded with a video camera. Alternation behavior was defined and counted as when mice entered all three of the arms consecutively (ABC, CAB, or BCA, but not BAB). The alternation count was divided by the maximum alternation (the total number of entries minus 2) and multiplied by 100 to create the percentage of alternation behavior. The total number of arm entries was recorded as spontaneous activity. Mice that entered arms fewer than 15 times during the test were excluded. None of the 30 mice used in this study exhibited total arm entries of fewer than 15 times.

### Rotarod test

In brief, mice were forced to run on a rotating rod with speeds starting at 5 r.p.m. and accelerating to 40 r.p.m. within 120 seconds and then remaining at 40 r.p.m. for 5 min. Mice were tested at 2 weeks, 3 weeks, and 4 weeks after surgery. In each session, three consecutive trials were conducted for each mouse with an interval of 15 minutes. The time a mouse stayed on the rotating rod was measured. Data were expressed as mean values from three trials.

### Elevated beam test

The elevated beam test was performed according to previous descriptions ^[56]^. The movement of mice on a round plastic beam (length 50 cm and diameter 1 cm) 40 cm above a surface with bedding was recorded and analyzed. The percentage of steps with hind paw slips during runs on the beam was calculated.

### TEM

TEM was used to measure myelin thickness in the corpus callosum area. Mice were anesthetized deeply and perfused intracardially with 2.5% glutaraldehyde in 4% paraformaldehyde (PFA). Brain tissue was cut into 1-mm slices using a mouse coronal slice mold and then extracted from the left corpus callosum approximately 1 mm^3^ from the site of 0.5 mm behind bregma. All specimens were post-fixed in osmium tetroxide, dehydrated through graded concentrations of ethanol, pre-embedded with propylene oxide, and flat-embedded in epoxy resin. Samples were processed for routine TEM observation and examined using an electron microscope (Hitachi, HT7700, Japan) at 120 kV. Three to five images were acquired in randomly selected areas within the corpus callosum at a magnification of 50,000× and analyzed with ImageJ by an investigator blinded to experimental groups. Over 50 axons per animal were analyzed by tracing the axonal circumference and the whole fiber circumference in a blinded fashion. G-ratios were determined as the ratio of the inner axonal diameter to the outer axonal diameter.

### LFB staining

LFB staining was used to detect any histological changes in white matter according to the manufacturer’s protocol. In brief, brain slices were incubated in 0.1% LFB (ab150675, Abcam) in acidified 95% ethanol overnight at 60 °C. The slices were then differentiated and counterstained with 0.05% lithium carbonate and cresyl violet solution. WMLs in the corpus callosum were observed under an Olympus SLIDEVIEW VS200 microscope (Invitrogen, USA) and quantitatively analyzed by Image J/Image pro-plus (Media Cybernetics, Silver Spring, USA).

### Immunohistochemistry, multiplex immunofluorescence staining, and image analysis

Immunostaining of brain slices was conducted as previously described ^[57]^. Primary antibodies were incubated at 4 °C overnight at vendor concentration. After washing with cold PBS five times, slices were incubated with fluorescence-conjugated secondary antibodies at room temperature for 1 hour. Slides were washed and mounted with a mounting medium containing 4’,6-diamidino-2-phenylindole (DAPI; ab104139, Abcam). Primary antibodies used were as follows: anti-mouse MBP (ab40390, 1:1,000, Abcam), anti-mouse Olig2 (1:500, AB9610, Millipore; 1:500, 211F1.1, MABN50, Millipore), anti-mouse APC/CC1 (ab16794, 1:200, Abcam), and anti-mouse NG2 (EPR23976-145, ab275024, 1:100, Abcam). Secondary antibodies used were as follows: Alexa Fluor 594 anti-rabbit IgG (A21207, Invitrogen), Alexa Fluor 594 anti-mouse IgG (A21203, Invitrogen), Alexa Fluor 488 anti-rabbit IgG (A21206, Invitrogen), and Alexa Fluor 488 anti-mouse IgG (A21202, Invitrogen). Human brain slices were incubated with anti-human ELANE (ab68672, 1:100, Abcam), anti-human Iba1 (EPR16589, ab283319, 1:300, Abcam) and anti-human CD66b (ab197678, 1:100, Abcam) antibodies to evaluate the co-expression of ELANE with microglia or neutrophil using an Opal 3-Plex Manual Detection kit (# NEL810001KT, Akoya Bioscience, Marlborough, MA, USA) following the manufacturer’s protocol. Images were collected with a Vectra 3.0 spectral imaging system (PerkinElmer, Waltham, MA) or a Zeiss LSM 710 confocal microscope (Carl Zeiss, Jena, Germany). Image analysis was performed using ImageJ software (National Institutes of Health, USA).

### Western blotting

Mice were euthanized by lethal anesthesia via isoflurane, and, after pericardiac perfusion with cold PBS, brain-dissected corpus callosum tissues and bone marrow cells from the femurs and tibias were collected for western blotting. Tissue samples or cell cultures were homogenized in radioimmunoprecipitation assay lysis buffer (Solarbio) with 1 mM phenyl-methanesulfonyl fluoride (Solarbio) and phosphatase inhibitor cocktails (sc-45065, Santa Cruz Biotechnology, Dallas, TX). After centrifugation, the supernatants were collected as total proteins. Then, the proteins were separated by 10% SDS-PAGE and transferred onto a polyvinylidene fluoride (PVDF) membrane (Amersham Biosciences). Immunoblot analysis was performed with primary antibodies against ELANE (E6K6Q, 61928, Cell Signaling Technology), MBP (D8X4Q, 78896, Cell Signaling Technology), and CNPase (D83E10, 5664, Cell Signaling Technology) at 4 °C overnight, and actin (13E5, 4970, Cell Signaling Technology) was used as an internal loading control. Then, the membrane was incubated with horseradish peroxidase (HRP)-conjugated secondary antibody for 1 hour at room temperature. Chemiluminescence signal was detected using a G:BOX chemiluminescence imaging system (Syngene, Cambridge, UK).

### Flow cytometry

Single-cell suspensions for flow cytometry from the brain and femur bones were prepared as previously reported ^[57]^. In brief, mice were euthanized by lethal anesthesia via isoflurane, and, after perfusion with cold PBS, brain and femur bones were removed. Brain tissues were minced and incubated with collagenase IV and deoxyribonuclease at 37 °C for 30 minutes. After removing the myelin debris by centrifugation in 30% Percoll, single cells were suspended in 1% bovine serum albumin followed by antibody staining. In experiments using bone marrow, single cells were collected after removal of red blood cells using lysis buffer. Single-cell suspensions were placed in conical centrifuge tubes (10^6^ cells per tube) and stained with fluorescently conjugated antibodies. For intracellular staining, cells were fixed in fixation buffer for 20 minutes after surface marker staining. After washing twice in permeabilization buffer, cells were incubated with antibodies in the staining buffer for 45 minutes. All flow cytometry antibodies were sourced from BioLegend, unless otherwise indicated. Antibodies used were as follows: CD11b-PE-Cy7 (M1/70, 101216), CD45-APC-Cy7 (30-F11, 103116), Zombie (423113), ELANE (E6K6Q, 61928, Cell Signaling Technology), and Neuropilin-1 (AF566, R&D Systems). Samples were run on a FACSAria (BD Biosciences) and analyzed using FACSDiva and FlowJo X software (Informer Technologies, Ashland, OR).

### ELISA for ELANE

The concentration of ELANE in the patient plasma samples was measured using a Human Neutrophil Elastase/ELA2 DuoSet ELISA kit (DY9167, R&D Systems). ELANE expression in mouse corpus callosum areas and plasma samples was measured using commercial ELISA kits (MELA20, R&D Systems). Mouse blood samples were obtained through a venous catheter placed in the inner canthus of the eye. Plasma samples were isolated from blood anticoagulated with EDTA by centrifugation (3,000*g* for 3 minutes) and were stored at −80 °C until used. Mice were euthanized by lethal anesthesia via isoflurane, and, after perfusion with cold PBS, the brain was removed. Dissected corpus callosum tissues were minced, and tissue homogenate supernatant samples were prepared by taking the top fraction after centrifugation at 1,0000*g* for 10 minutes at 4°C. Total protein concentration from these tissue homogenate supernatant samples was determined using a BCA protein assay kit (Thermo Fisher Scientific). In brief, supernatant fluid was added to separate microplates containing specific antibodies. Enzyme conjugate was dispensed into each well, and the plate was incubated at 37 °C for 60 minutes. Then, the plate was washed five times. Subsequently, Chromogen A and Chromogen B reagent were dispensed into each well, and the plate was incubated at room temperature in the dark for 20 minutes. Finally, the reaction was stopped by adding Stop Solution, and the absorbance at 450 nm was determined using a microplate reader.

### ELANE activity assays

ELANE catalytic activity was determined using the synthetic substrate *N*-methoxysuccinyl-Ala-Ala-Pro-Val p-nitroanilide (M4765, Sigma-Aldrich, St. Louis, MO). In brief, plasma or corpus callosum tissue homogenate samples were incubated in 0.1 M Tris-HCl buffer (pH 8.0) containing 0.5 M NaCl and 1 mM substrate for 24 hours at 37 °C. After incubation, p-nitroanilide release was measured spectrophotometrically at 405 nm and was considered as ELANE activity. For inactivation, ELANE was incubated with PMSF (329-98-6, Sigma-Aldrich) for 2 hours, and residual PMSF was eliminated with a PD-10 desalting column (GE Healthcare Life Sciences, Uppsala, Sweden).

### Mass spectrometry to determine CNPase cleavage sites

ELANE cleavage sites on CNPase were determined by incubating recombinant human CNPase (aa 1–421, MBS1475160, MyBioSource, 4 μg) with human native ELANE (SE563, Elastin Products Company, 0.25 μg) for 2 hours at 37 °C, and reactions were stopped by boiling in SDS loading buffer. The cleavage products were subjected to N-terminal dimethyl labeling and analyzed using liquid chromatography–tandem mass spectrometry (LC–MS/MS) at Beijing Bio-tech Pack Technology Co., Ltd. (Beijing, China). In brief, protein dimethyl labeling was performed on protein fragments by the addition of 40 nM formaldehyde (ultrapure grade) in the presence of 20 nM sodium cyanoborohydride and incubated at 37 °C for 1 hour. The reaction was quenched by the addition of 100 mM ammonium bicarbonate and precipitated with 8 volumes of acetone and 1 volume of methanol at 20 °C for 3 hours. The precipitated protein underwent reductive alkylation with 10 mM dithiothreitol (Sigma-Aldrich) at 56 °C for 1 hour and was alkylated by 55 mM iodoacetamide at room temperature in the dark for 40 minutes. Subsequently, enzymatic digestion using either trypsin or chymotrypsin was performed, and the resulting peptides were subjected to LC–MS/MS analysis using an Orbitrap Fusion Lumos Tribrid mass spectrometer coupled with a Vanquish UHPLC system (Thermo Fisher Scientific, USA). The sequences were searched in the UniProt human CNPase database (P09543) using Byonic (Protein Metrics, San Carlos, CA).

### Tissue dissociation and cell isolation by magnetic cell separation

Murine neonatal cerebellar tissue from newborn (P5–P7) C57BL/6N mice, of either sex, was dissected, weighed and enzymatically digested using a Neural Tissue Dissociation Kit (130-092-628 for microglia and oligodendrocyte progenitors and neurons; 130-093-231 for astrocytes; Miltenyi Biotec, Germany Germany). Dissociated tissue was passed through a 100-μm cell strainer (BD Bioscience) and pelleted by 10-minute centrifugation at 300*g* at room temperature. The cell pellet was resuspended in D-PBS buffer and processed immediately.

Magnetic-activated cell sorting (MACS) was used to specifically enrich different neural cells in murine brain cell populations. All microbead kits were from Miltenyi Biotec, unless otherwise indicated. Microglia, astrocytes, and OPCs were magnetically labeled with anti-CD11b microbeads (130-093-636), anti-GLAST (ACSA-1) microbeads (130-095-825), and anti-CD140a (PDGFRα) microbeads (130-101-502), respectively, and passed through a MACS column, which was placed in the magnetic field of a MACS Separator (Miltenyi Biotec, Germany). Neurons were isolated by negative selection using a mouse neuron isolation kit (130-115-390). The amounts of antibodies and magnetic beads were calculated based on the number of cells obtained after myelin removal (130-096-733) according to the manufacturer’s instructions. The sorted cells were verified by immunofluorescence staining with antibodies to cell-specific markers.

### Primary culture and treatment of mouse OPCs and differentiated oligodendrocytes

Primary OPCs were obtained from P5–P7 C57BL/6N pups using MACS with the CD140a (PDGFRα) MicroBead Kit (130-101-502, Miltenyi Biotec, Germany) according to the manufacturer’s protocol. Primary OPCs were plated into poly-D-lysine/laminin-coated coverslips and expanded in neurobasal medium (A2477501, Gibco) with 2% B27 (17504044, Invitrogen, USA), 10 ng/ml PDGF-AA (100-13A, PeproTech, NJ, USA), 10 ng/mL CNTF (450-50, PeproTech), and 1 ng/ml NT3 (450-03, PeproTech) for 3 days before experiments. For OPC differentiation, the medium was completely replaced with the differentiation medium. Differentiation medium components included 2% B27 supplement, 5 μg/ml insulin (I2643, Sigma-Aldrich), 40 ng/ml triiodo-thyronine (T3) (T6397, Sigma-Aldrich), and 40 ng/ml CNTF in neurobasal medium. OPCs were allowed to differentiate in differentiation medium for 4–8 days with one half medium changed every 2– 3 days. Human native ELANE was labeled with FITC according to the protocol supplied by the ReadiLink Rapid FITC Antibody Labeling Kit (AAT Bioquest Inc., USA). For inhibition of endocytosis, mouse OPCs were pre-incubated with the endocytosis inhibitor Dynasore (D7693, Sigma-Aldrich) before ELANE treatment.

### CG4 cell culture and transfection

CG4 cells were originally derived from rat glial progenitor cells that were induced to differentiate into myelin-producing cells ^[58]^. This cell line is widely used to study the biology of oligodendrocyte progenitor cells. CG4 cells were cultured on poly-D-lysine-coated tissue culture dishes in growth medium containing DMEM-F12 (11330032, Gibco), 2% B27 supplement (17504044, Invitrogen), 1% N_2_ supplement (17502048, Invitrogen), 10 ng/ml PDGF-AA (100-13A, PeproTech), and 10 ng/ml EGF (315-09, PeproTech). CG4 cells were transfected with a full-length human *CNPase* construct in pcDNA3.1 vector (Invitrogen) or with an empty pcDNA3.1 vector, using Lipofectamine 2000 transfection reagent (Invitrogen) according to the manufacturer’s protocol. The small interfering RNA (siRNA) sequences were obtained from GenePharma (Shanghai, China), including a scrambled siRNA (silencer negative control) sequence and two CNPase siRNA sequences. The target sequences are shown in Table S2. siRNA transfection was performed with Lipofectamine RNAiMAX (Invitrogen) in serum-free/antibiotic-free medium. After a 48-hour period, cells were collected, and expression of CNPase was confirmed by western blot.

### HEK293T cell culture, plasmid construction, and transfection

To create CNPase mutants that were refractory to cleavage by ELANE, we replaced all residues at P1 and P2 positions in all 11 cleavage sites with tryptophan residues. The full-length coding sequences of WT CNPase, mutated CNPases, and ELANE were synthesized commercially by SynbioB Inc. (Tianjin, China). The synthesized nucleotides were cloned into the mammalian expression vector pcDNA3.1 (Invitrogen, Carlsbad, CA, USA) under the cytomegalovirus (CMV) promotor according to standard techniques. The recombinant plasmids were designated as pcDNA3.1/CNPase-wt, pcDNA3.1/CNPase-mut, and pcDNA3.1/ELANE, respectively, and were verified by nucleotide sequencing.

HEK293T cells were maintained in DMEM medium (11965092, Gibco) supplemented with 10% fetal bovine serum (FBS) (26140079, Gibco), 100 U/ml penicillin and 100 mg/ml streptomycin, maintained at 37 °C with 5% CO_2_. The plasmid pcDNA3.1/CNPase-wt or pcDNA3.1/CNPase-mut was transiently co-transfected with plasmid pcDNA3.1/ELANE into HEK293 cells. In brief, cells were seeded in six-well plates and allowed to grow for 24 hours until reaching 50%–60% confluence. Transfection complexes were prepared in serum-free Opti-MEM media in a ratio Lipo:DNA (1:1). Mixtures were incubated for 10 minutes at room temperature and then added drop-wise to the cells in serum medium. After 5 hours, transfection medium was replaced with fresh complete DMEM, and cells were returned to the incubator for an additional 48 hours.

### Cell viability assays

Cell viability was assayed using the CellTiter-Glo luminescent cell viability assay kit (G7570, Promega) according to the manufacturer’s instructions. In brief, cells were plated and treated at the time points indicated in triplicate for each experimental condition. After treatment, 100 μl of CellTiter-Glo reagent was added to each well. The contents were mixed for 2 minutes to induce cellular lysis followed by incubation at room temperature for 10 minutes to stabilize the signal. Then, the luminescence derived from the luciferase reaction was immediately recorded on a Microplate Reader (Bio-TEK Instruments, Winooski, Vt).

### Immunofluorescence

Primary oligodendrocytes were isolated as described above and grown on poly-D-lysine-coated glass slides. At the indicated time point, the cells were washed with PBS before fixation with 4% paraformaldehyde (in PBS, pH 7.4) for 15 minutes at room temperature. The cells were permeabilized with 0.1% Triton X-100 for 15 minutes, blocked with 5% BSA/PBS for 1 hour, and incubated with primary antibodies at 4 °C overnight. After washing with PBS, the cells were incubated with fluorochrome-conjugated secondary antibodies (1:1,000 dilution, Invitrogen) for 1 hour at room temperature. Finally, nuclei were counterstained with DAPI. Primary antibodies used were as follows: anti-mouse A2B5 (ab53521, Abcam), anti-mouse NG2 (EPR23976-145, ab275024, Abcam), anti-mouse MBP (ab40390, Abcam), anti-mouse CNPase (ab6319, Abcam), and anti-mouse cleaved caspase-3 (5A1E, 9664, Cell Signaling Technology). All images were acquired on a Zeiss LSM 710 confocal microscope (Carl Zeiss, Jena, Germany). Image analysis was performed using ImageJ software (National Institutes of Health, USA).

### Statistical analysis

Categorical variables within clinical data are presented as frequencies and percentages. Continuous variables with skewed distributions are expressed as median and interquartile range. Patients were categorized into tertiles based on plasma ELANE levels. The nonparametric Wilcoxon test or Kruskal-Wallis test was used to analyze differences in continuous variables, and the *X*^2^ test was employed to analyze differences in categorical variables.

Multivariate logistic regression models and generalized linear models were used to explore the relationship between ELANE levels and adverse outcomes, using the lowest ELANE tertile group as the reference group. Variables were included in the adjustment if they were associated with ELANE levels or if they were traditional predictors of recurrent stroke. Unadjusted and adjusted relative risks (RRs), odds ratios (ORs), and their 95% confidence intervals (CIs) were calculated. Additionally, Pearson’s correlation coefficient was applied to compute r values and p values, which were used to measure the linear correlation between plasma ELANE levels and adverse outcomes.

All data from *in vitro* and *in vivo* experiments in text and figures are presented as the mean ± s.e.m. The Shapiro–Wilk normality test was used to confirm the values derived from a Gaussian distribution. Assumptions of equal variance were tested with Brown–Forsythe tests. Statistical significance was determined by two-tailed unpaired Student’s *t*-test for two groups, Mann–Whitney test for non-parametric data, one-way ANOVA followed by Tukey’s post hoc test for three or more groups, or two-way ANOVA accompanied by Bonferroni post hoc test for multiple comparisons. The criterion of significance was *P* < 0.05. All statistical analyses were performed using Prism 8.0 software (GraphPad Software, San Diego, CA, USA).

## Notes

### Competing Interest Statement

The authors have declared no competing interest.

